# Reference-based assignment of aerolysin nanopore signals to peptide sequences from tryptic protein digests

**DOI:** 10.64898/2026.10.21.760613

**Authors:** Carolin Hensel, Carolin Kleber, Ida Suppanz, Atefeh Akbari, Sebastian Gutsch, Tobias Hutzenlaub, Tobias Ensslen, Jan C. Behrends, Felix von Stetten, Gerhard Mittler, Jan-Niklas Klatt, Andreas Schreiber

## Abstract

Rapid, cost-efficient and miniaturized protein analysis is essential for point-of-care clinical diagnostics. Nanopores offer a promising alternative to mass spectrometry by enabling label-free, single-molecule sensing with a minimal footprint. However, peptide-level protein identification remains challenging because of the difficulty of assigning nanopore signals to peptide sequences and the incompatibility of conventional proteolysis workflows with nanopore sensing. Here, we show a reference-based strategy for peptide assignment in protein digests by combining nanopore measurements of synthetic peptides with LC-MS/MS analysis. Tryptic digests of myelin basic protein (MBP), prepared using a single-pot solid-phase-enhanced (SP3) workflow to remove nanopore-incompatible sample preparation reagents, yielded a reproducible five-population fingerprint using an engineered aerolysin nanopore (AeL-R220S). Comparison with synthetic peptide references and cross-validation with LC-MS/MS enabled the assignment of all five blockade populations to four MBP-derived peptides, although overlapping blockade distributions limited the resolution of peptides with comparable molecular volumes. This approach establishes a reference-based assignment strategy and compatible sample-preparation workflow for nanopore sensing, paving the way for label-free identification and characterization of protein targets.

## Introduction

The advancement of clinical diagnostics and biomedical research has created a growing demand for analytical platforms capable of rapid, cost-efficient and miniaturized protein analysis. Liquid chromatography tandem mass spectrometry (LC-MS/MS) remains the gold standard for proteomic profiling due to its high versatility in accommodating a broad range of analyte sizes with distinct chemical properties while providing high accuracy and sensitivity even within complex biological matrices.^1^ However, LC-MS/MS requires spatially demanding and maintenance-intensive high-end instrumentation operated by trained personnel, which restricts LC-MS/MS to centralized facilities and hinders its deployment in point-of-care (POC) settings.^2^

Nanopore sensing has emerged as a compelling, low-footprint alternative, combining single-molecule sensitivity and real-time readout with low operational costs and portable instrumentation.^3–6^ In biological nanopore sensing, an applied transmembrane potential drives solute ions through a single pore embedded in a synthetic lipid bilayer. As individual analyte molecules enter the pore lumen, they partially occlude the ionic current, generating characteristic residual current blockades (*I/I*_*o*_) and dwell times (*τ*) that reflect analyte volume and conformational details.^7^ Numerous studies have demonstrated single-molecule sensing with monomer resolution^8–13^, successfully resolving subtle molecular differences such as post-translational modifications (PTMs)^13–17^ and even isomerization.^13,14,18–21^. Despite this formidable resolution, direct nanopore analysis of full-length proteins still remains elusive. While a few studies have demonstrated detection of entire, small native proteins,^22–24^ larger folded targets are spatially demanding and frequently exceed the pore’s internal volume, whereas chemical denaturants used to unfold large proteins often induce heterogeneous translocation behavior.^25–27^ Although active translocation using motor proteins to processively feed unfolded polypeptide chains through a pore offers a promising workaround, this strategy remains constrained by stochastic stepping dynamics of the motor proteins.^28,29^ Recent advances have further expanded the scope of nanopore-based protein analysis, including DNA-assisted, parallel peptide profiling^30^ and high-resolution peptide sensing using engineered MspA nanopores^31^.

Mirroring bottom-up mass spectrometry strategies, proteolysis breaks intact proteins into smaller peptide fragments, compatible with nanopores to generate characteristic peptide fingerprints that enable protein identification through unique patterns of current blockades.^32,33^ Specifically, Lucas et al.^32^ demonstrated that nanopore peptide profiling can enable protein identification and Bakshloo et al.^33^ further advanced nanopore-based protein identification *in situ* using fragment-based readout via tryptic peptides. However, direct assignment of nanopore signals to specific peptide sequences has remained challenging.

A major hurdle for fragmentation-based nanopore analysis is the limited compatibility of sample preparation workflows with nanopore sensing. Conventional protein digestion workflows require reagents such as sodium dodecyl sulfate (SDS) for protein denaturation^34,35^ and reducing agents such as DTT (dithiothreitol) or TCEP (tris(2-carboxyethyl)phosphine).^36^ These reagents, although essential for efficient sample processing, compromise the stability of lipid bilayers^37,38^, biological nanopores or Ag/AgCl electrodes^39^, hindering direct adoption of established proteomic workflows for nanopore analysis. Consistent with the mentioned membrane-disrupting effects of detergents, our preliminary nanopore measurements of unpurified standard in-solution digests of proteins (see Supplementary Note 1) led to rapid membrane destabilization and loss of stable open-pore currents, highlighting the need for efficient removal of nanopore-incompatible reagents.

Importantly, detergents and other sample preparation reagents can also interfere with electrospray ionization in LC-MS/MS, and established proteomic workflows therefore already incorporate dedicated cleanup strategies to remove these compounds.^40^ These established approaches provide a promising starting point for adapting sample purification to nanopore-based protein analysis. However, whether the resulting degree of purification is sufficient to ensure stable nanopore operation and reliable peptide detection remains to be experimentally evaluated.

Beyond reagent compatibility, the resulting peptides must meet the physicochemical requirements of the nanopore assay. Under idealized experimental conditions using the aerolysin (AeL) nanopore, peptides are typically required at nano-to micromolar concentrations and within a length range of approximately 4-20 amino acids to ensure sufficient entrapment within the pore for reliable determination of residual current levels.^7^ The single-pot, solid-phase-enhanced sample preparation (SP3) workflow,^40^ already established in LC-MS/MS proteomics, provides compatibility with nanopore sensing conditions and therefore represents a highly attractive sample preparation protocol. The SP3 workflow integrates enzymatic proteolysis with efficient removal of nanopore-incompatible reagents by a magnetic bead-based cleanup within a single tube.

As a proof of concept, we demonstrate the applicability of the SP3 workflow using bovine-derived myelin basic protein (MBP) as a clinically relevant model protein. MBP is a major component of the myelin sheath and its breakdown products in cerebrospinal fluid or serum serve as important biomarkers of multiple sclerosis (MS) activity and exacerbations, while also informing prognosis and being associated with cognitive dysfunction.^41^

Following SP3 tryptic digestion of MBP, the resulting peptides were analyzed using an engineered aerolysin nanopore (AeL-R220S)^14,42^, with optimized sensitivity for positively charged peptides. The SP3-based workflow renders proteolytic digests compatible with nanopore sensing and demonstrates, through cross-validation with LC-MS/MS, the reference-based assignment of five predominant nanopore blockade signatures to four specific peptide identities.

## Results & Discussion

### SP3-processing of myelin basic protein (MBP)

Myelin basic protein was processed using the established single-pot solid-phase-enhanced (SP3) workflow, adapted here to yield a digest that can be measured directly by nanopore sensing without further purification (Fig. 1). Experimental digestion conditions were optimized by evaluation of LC-MS/MS peptide intensities, benchmarked against an *in silico* tryptic digestion model accounting for site-specific cleavage probabilities (Supplementary Fig. 1). LC-MS/MS analysis showed that increasing the SDS concentration from 0 to 1% (w/v) prior to bead binding resulted in complete sequence coverage (Supplementary Fig. 2) and increased the relative MS signal attributed to fully cleaved peptides, which accounted for 97.9% of the total peptide intensity (Supplementary Fig. 3), indicating improved digestion efficiency. For comprehensive LC-MS/MS results see Supplementary Table 1.

**Fig. 1:**
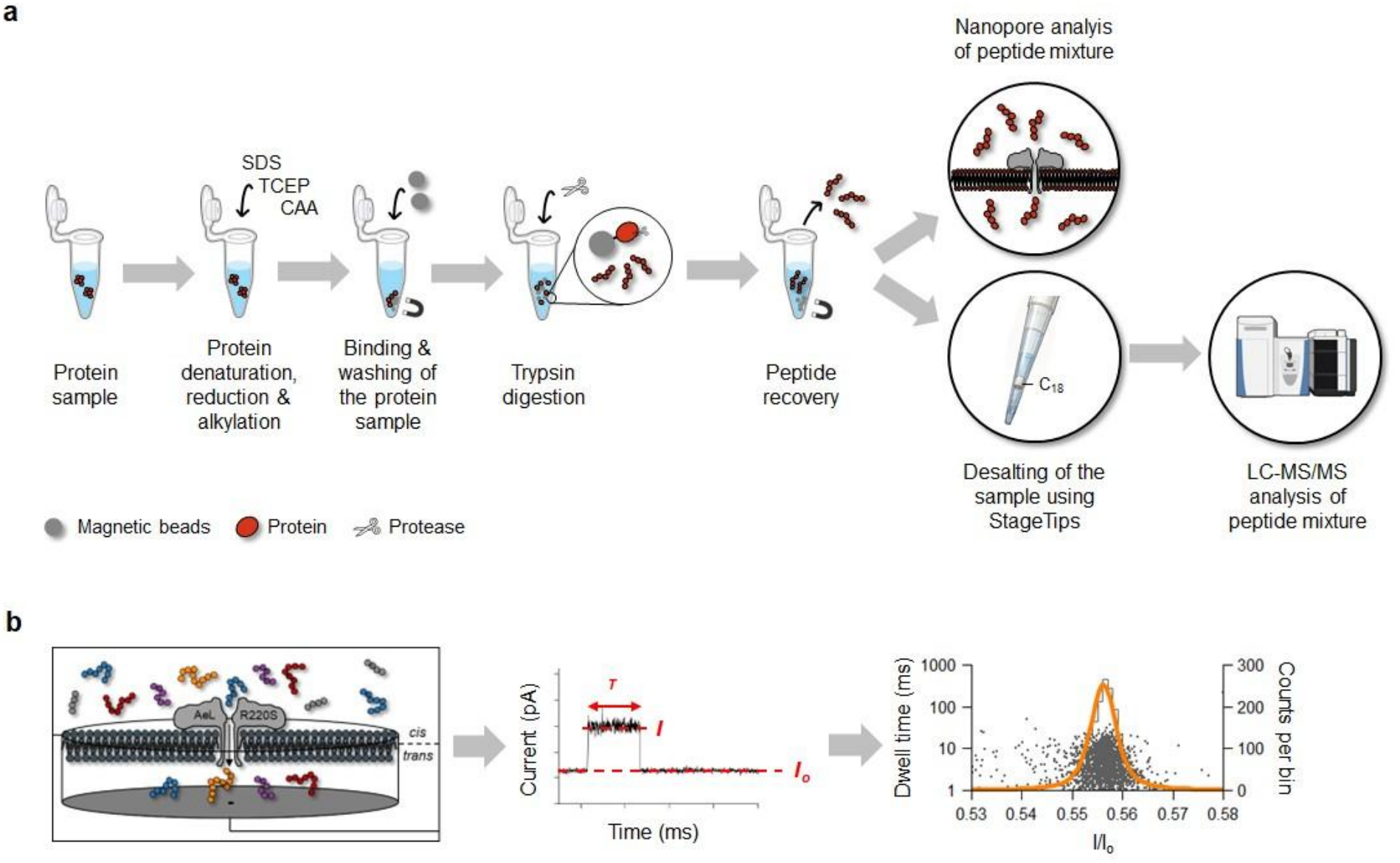
SP3 workflow for parallel nanopore sensing and LC-MS/MS analysis. **a** Proteins are denatured (SDS), followed by disulfide-bond reduction (TCEP) and alkylation (CAA) prior to adsorption onto carboxylate-functionalized magnetic beads. Subsequent washing eliminates nanopore-incompatible reagents, followed by on-bead tryptic proteolysis. Eluted peptides can be analyzed by both, nanopore sensing and, with an additional C_18_-based desalting step, LC-MS/MS analysis. **b** Nanopore analysis of peptides. Left: Schematic of the AeL-R220S nanopore reconstituted in a lipid bilayer, with peptides captured from *cis* to *trans*. Center: Analyte trapping in the pore produces transient current blockades, which are detected from the recorded current trace using a sliding-window baseline correction followed by threshold-based event detection. For each detected event, the relative residual current blockades (*I/I*_*o*_) and exponentially distributed dwell times (*τ*) are determined. Events with dwell times < 1 ms are excluded from subsequent analysis because their durations cannot be reliably resolved under the applied recording conditions. Right: Detected blockade events are displayed as scatter plots of *I/I*_*o*_ versus *τ* (grey). The *I/I*_*o*_ values are subsequently binned into histograms (black; bin width = 0.001) and Voigt functions^43^ (orange) are fitted to the resulting distributions to determine the peak maxima.

Additionally, SP3-based magnetic bead cleanup enabled efficient removal of residual detergent to allow direct downstream analysis of the sample by nanopores. Notably, the resulting SP3-processed digest was applied directly to the nanopore measurement chamber without additional purification, while parallel LC-MS/MS analysis required an additional desalting step to facilitate effective electrospray ionization.

### Nanopore analysis of SP3-processed MBP digest

Nanopore measurements of the SP3-processed MBP digest were performed using the Aerolysin (AeL) variant AeL-R220S^14,42^. Transmembrane potentials from -40 mV to -90 mV were screened and -60 mV was selected for all subsequent measurements as it gave the clearest separation between event populations (Supplementary Fig. 5). Interrogating the SP3-derived MBP digest at a nominal final concentration of 1 µM revealed a characteristic peptide fingerprint comprising five distinct blockade amplitudes (Fig. 2a-c). Across independent nanopore measurements (n = 3), the fingerprint revealed reproducible *I/I*_*o*_ values with a standard deviation of ≤ 0.004, corresponding to a relative standard deviation of ≤ 0.59 % (Supplementary Table 2, Supplementary Fig. 6). Peaks 2-5 were fitted in all three measurements. Events in the *I/I*_*o*_ range of peak 1 occurred in all three recordings, but in one measurement only three such events were detected, which was insufficient for a Voigt fit; peak 1 was therefore fitted in two of three measurements. Event frequencies for every peak are given in Supplementary Table 2. This reflects the low abundance of species 1 in the digest together with a low capture efficiency, which complicates its detection by nanopore sensing.

**Fig. 2:**
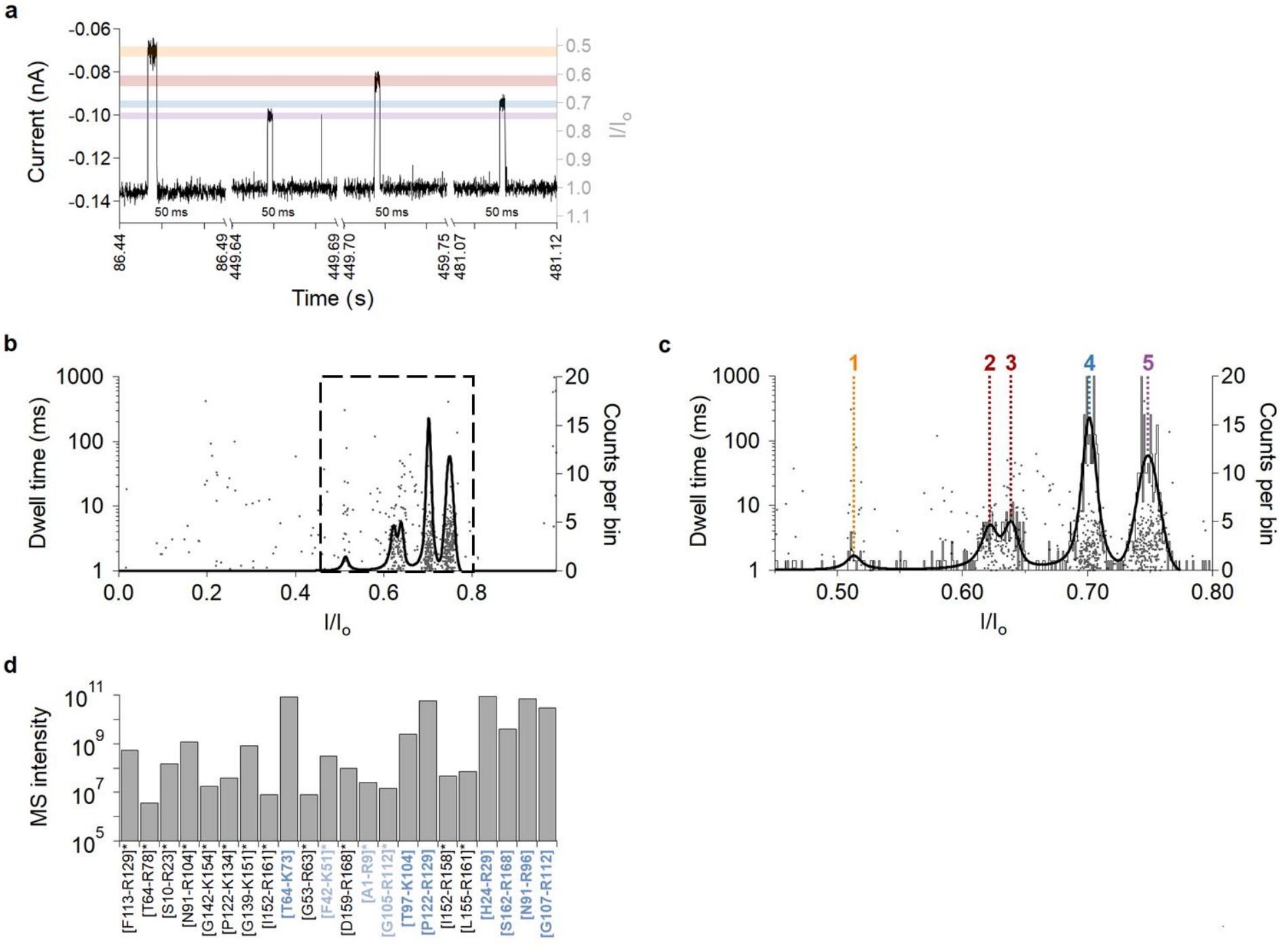
Single-molecule nanopore fingerprinting and LC-MS/MS characterization of SP3-processed MBP digest. **a** Representative fragment of ionic current recording of the MBP digest. **b-c** Scatter plot (grey) of the dwell time versus the relative residual current (*I/I*_*o*_) of the MBP digest (nominal final concentration: 1 µM) recorded with AeL-R220S at an applied potential of -60 mV. Voigt functions were fitted to histograms of the corresponding *I/I*_*o*_ distributions yielding the overall fitting profile (black).^43^ The dataset comprise independent nanopore measurements (n = 3) of the SP3-processed MBP digest combined into a single dataset. **b** Full *I/I*_*o*_ range. The dashed line indicates the range magnified in c. **c** Magnified view of the *I/I*_*o*_ range containing the five-population fingerprint observed for the MBP digest. **d** LC-MS/MS profiling of the MBP digest serving as reference library. Measured peptide intensities are plotted for candidates meeting nanopore compatibility criteria (length: 4-20 amino acids; net positive charge), ordered by theoretical peptide volume (decreasing from left to right). Volumes were calculated using the Peptide Property Calculator.^44^ Asterisks (*) denote peptides containing missed cleavages. Peptides highlighted in the figure were selected for synthetic synthesis to establish peptide assignment.

Importantly, parallel LC-MS/MS analysis of the SP3-processed MBP digest provided a reference for nanopore analysis. The identified peptides covered the MBP sequence (Supplementary Table 1), with identities and relative intensities of candidates meeting nanopore compatibility criteria visualized in Fig. 2d To further ensure that the observed nanopore events indeed originated from MBP-derived peptides rather than from trypsin autolysis products, a trypsin control was performed (see Supplementary Note 2). Nanopore analysis of the trypsin control resulted in only rare events (Supplementary Fig. 7) and their blockade amplitudes did not correspond to the MBP peptide fingerprint.

Altogether, these results demonstrate that the SP3 workflow generates tryptic peptides of appropriate length, concentration and purity for direct nanopore sensing, successfully overcoming the limitations caused by chemical interference typical of standard in-solution digestion protocols (see Supplementary Note 1).

### Nanopore analysis of synthetic MBP peptides

Beyond fingerprint detection,^32,33^ achieving assignment of individual blockade populations to specific peptides is essential for reliable protein identification. In gold-standard bottom-up mass spectrometry, peptide identification is achieved by matching precursor and fragment ion spectra against reference protein databases, with protein identification typically based on multiple unique peptides and controlled false discovery rates.^45^ As equivalent reference databases are not yet available for nanopore measurements, we instead measured individual synthetic MBP peptides to establish a basis for assigning specific blockade populations to their corresponding peptide sequences.

Based on the *in silico* digestion of MBP (Supplementary Fig. 1), seven peptides were selected based on their fulfillment of the physicochemical requirements for nanopore measurements, including a peptide length of 4-20 amino acids and a positive net charge. Importantly, all seven peptides were also detected by LC-MS/MS, confirming their presence in the MBP digest. These seven peptides were subsequently synthesized (Supplementary Table 3) and individually characterized using the AeL-R220S nanopore to establish a reference dataset of their characteristic blockade signatures.

At a peptide concentration of 5 µM, five of the seven selected peptides could be successfully detected using the AeL-R220S nanopore (Fig. 3a, Table 1). The remaining two peptides ([S162–R168] and [G107–R112]) generated only a few blockade events under the applied measurement conditions, insufficient to establish statistically meaningful blockade distributions or perform reliable peak fitting. Notably, both peptides have a calculated net charge of +0.76 at pH 7.5 (Table 1), comparable to several successfully detected peptides ([T97-R104], [P122-R129] and [N91-R96]). Thus, despite fulfilling the applied physicochemical selection criteria and being confirmed as present in the MBP digest by LC-MS/MS, these peptides remained undetectable even at a final concentration of 5 µM, which exceeded their estimated concentrations in the digest. Triplicate measurements of the five detectable peptides demonstrated high reproducibility, with relative standard deviations (RSD) of the relative residual current values of ≤ 1.50% and corresponding standard deviations (SD) of 0.007 across independent experiments (Table 1, Supplementary Fig. 8).

**Fig. 3:**
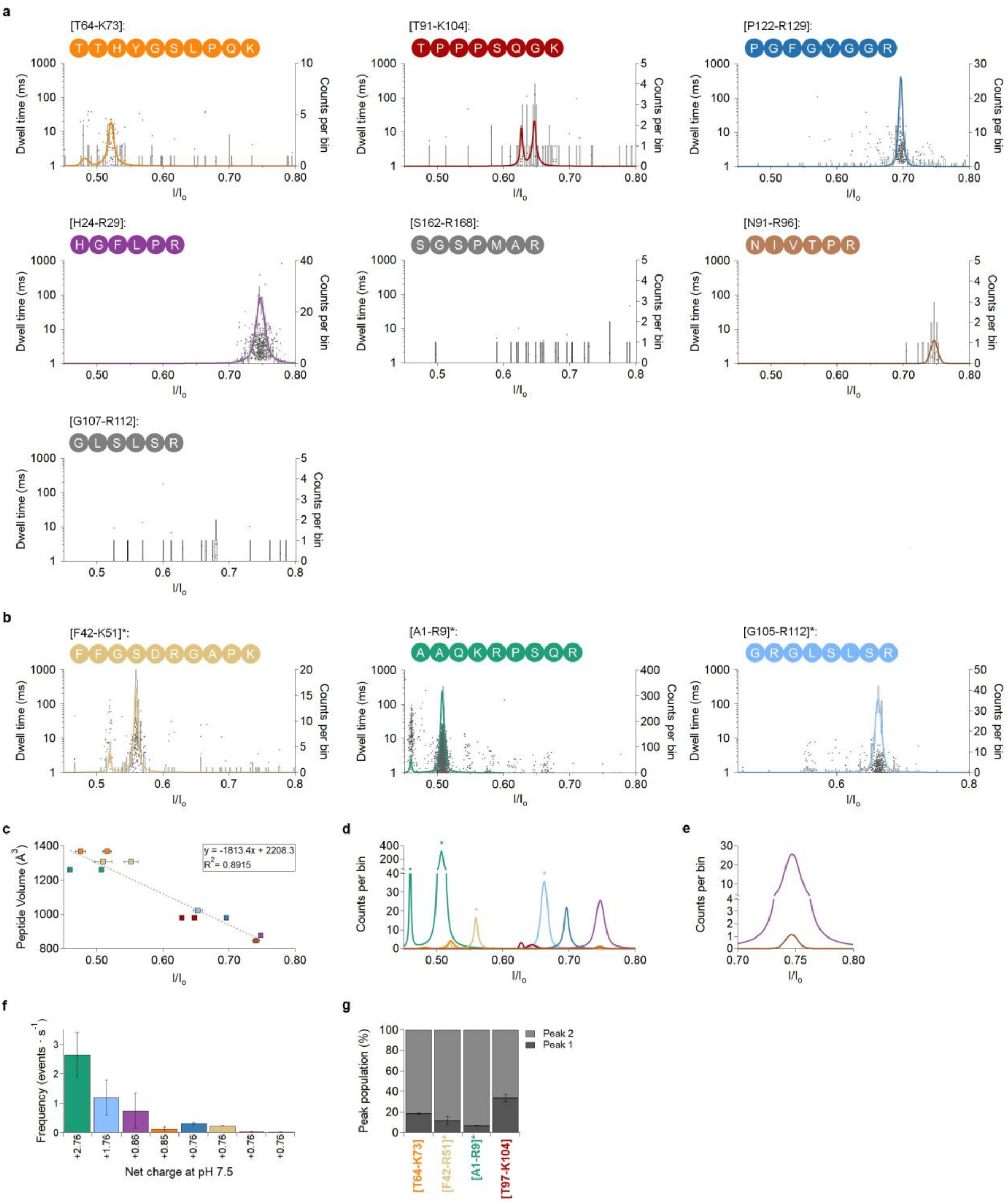
Nanopore analysis of selected synthetic MBP reference peptides. **a-b** Scatter plots (grey) of the dwell time versus the relative residual current (*I/I*_*o*_) for synthetic MBP peptides (nominal final concentration: 5 μM) measured with AeL-R220S at -60 mV. Voigt functions^43^ (colored) were fitted to histograms of the corresponding *I/I*_*o*_ distributions. Data were obtained from three independent measurements (n = 3), with representative datasets shown. **a** Nanopore analysis of peptides without missed cleavages, predicted by *in silico* digestion and confirmed by LC-MS/MS. Two peptides ([S162-R168] and [G105-R112]) were not detectable under the applied conditions; therefore, no Voigt fits are shown for these peptides. Peptides are ordered by their theoretical peptide volume (decreasing from left to right). **b** Nanopore analysis of peptides containing missed cleavages (marked with an asterisks), which were additionally identified by LC-MS/MS and included to provide further information for the assignment of the nanopore fingerprint. **c** Linear regression of detected *I/I*_*o*_ values against calculated molecular volumes (Å^3^). Peptides exhibiting two distinct blockade populations are represented by two data points at the same molecular volume. Error bars indicate the standard deviation (SD) of three independent measurements (n = 3). **d** Combined representation of the Voigt-fitted *I/I*_*o*_ distributions of all detectable synthetic MBP peptides shown in a and b. **e** Combined representation of the Voigt-fitted *I/I*_*o*_ distributions of [H24-R29] and [N91-R96] illustrating their substantial overlap, which is consistent with their similar molecular volumes. **f** Event frequencies of the measured peptides as a function of their calculated net charge at pH 7.5, showing an association between increased net charge and increased event frequency. Event frequencies were calculated by dividing the number of detected events by the measurement duration (~ 600 s). Error bars indicate the standard deviation (SD) of three independent measurements (n = 3). **g** Relative contribution of the two peak populations for peptides exhibiting bimodal blockade distributions, with the minor population accounting for 7-34% of the total events.

**Table 1:** Synthetic MBP peptides used for nanopore reference measurements, their physicochemical properties, and corresponding nanopore characteristics. Peptides are ordered by their theoretical peptide volume (decreasing from top to bottom). Peptide volumes were calculated using the Peptide Property Calculator.^44^ The theoretical net charge at pH 7.5 was calculated using the Henderson-Hasselbalch equation implemented in the MolDraw Peptide Calculator,^48^ using the pKa values reported by Thurlkill et al.^49^ The ionization states of the N- and C-termini and ionizable amino acid side chains were taken into account. The normalized residual current (*I/I*_*o*_), full width at half maximum (FWHM) and event frequency (at f.c. 5 µM) are reported as mean ± standard deviation (SD) from three independent measurements (n = 3), performed at an applied voltage of -60 mV. Measurements were recorded for approximately 600 s. For some peptides, two distinct current populations (Peak 1 and Peak 2) were observed, resulting in two corresponding *I/I*_*o*_ and event frequency values. Asterisks (*) indicate peptides containing missed cleavage sites.

| **Peptide** | **Sequence** | **Volume** (Å<sup>3</sup>) | **Net charge (pH 7.5)** | ***I/I<sub>o</sub>*** | **FWHM** | **Frequency (events·s<sup>-1</sup>)** |
|---|---|---|---|---|---|---|
| **[T64-K73]** | TTHYGSLPQK | 1369 | + 0.85 | **Peak 1:**<br>0.476 ± 0.007<br>**Peak 2:**<br>0.516 ± 0.006 | **Peak 1:**<br>0.0092 ± 0.0024<br>**Peak 2:**<br>0.0091 ± 0.0015 | **Peak 1:**<br>0.025 ± 0.014<br>**Peak 2:**<br>0.108 ± 0.051 |
| **[F42-K51]\*** | FFGSDRGAPK | 1308 | + 0.76 | **Peak 1:**<br>0.510 ± 0.013<br>**Peak 2:**<br>0.552 ± 0.010 | **Peak 1:**<br>0.0039 ± 0.0020<br>**Peak 2:**<br>0.0075 ± 0.0027 | **Peak 1:**<br>0.025 ± 0.009<br>Peak 2:<br>0.191 ± 0.008 |
| **[A1-R9]\*** | AAQKRPSQR | 1260 | + 2.76 | **Peak 1:**<br>0.460 ± 0.002<br>**Peak 2:**<br>0.507 ± 0.002 | **Peak 1:**<br>0.0022 ± 0.0003<br>**Peak 2:**<br>0.0063 ± 0.0005 | **Peak 1:**<br>0.175 ± 0.060<br>**Peak 2:**<br>2.47 ± 0.69 |
| **[G105-R112]\*** | GRGLSLSR | 1022 | + 1.76 | 0.653 ± 0.007 | 0.0105 ± 0.0014 | 1.19 ± 0.61 |
| **[T97-K104]** | TPPPSQGK | 981 | + 0.76 | **Peak 1:**<br>0.629 ± 0.002<br>**Peak 2:**<br>0.648 ± 0.004 | **Peak 1:**<br>0.0043 ± 0.0009<br>**Peak 2:**<br>0.0083 ± 0.0057 | **Peak 1:**<br>0.014 ± 0.002<br>**Peak 2:**<br>0.027 ± 0.003 |
| **[P122-R129]** | PGFGYGGR | 980 | + 0.76 | 0.696 ± 0.001 | 0.0093 ± 0.0020 | 0.313 ± 0.052 |
| **[H24-R29]** | HGFLPR | 878 | + 0.86 | 0.748 ± 0.003 | 0.0114 ± 0.0035 | 0.757 ± 0.034 |
| **[S162-R168]** | SGSPMAR | 853 | + 0.76 | Not detectable |  |  |
| **[N91-R96]** | NIVTPR | 846 | + 0.76 | 0.741 ± 0.006 | 0.0089 ± 0.0026 | 0.019 ± 0.005 |
| **[G107-R112]** | GLSLSR | 765 | + 0.76 | Not detectable |  |  |

Beyond the fully cleaved peptides predicted by *in silico* digestion, LC-MS/MS also identified several peptides containing missed cleavages (Fig. 2d, Supplementary Table 1). These peptides generally exhibited lower MS/MS signal intensities than their fully cleaved counterparts, potentially reflecting differences in peptide abundance and/or ionization efficiency.

To further support the nanopore peak assignment, three representative missed cleavage peptides ([F42-K51]*, [A1-R9]*, and [G105-R112]*) were selected for additional nanopore measurements. The selection focused on peptides with molecular volumes comparable to those of already characterized fully cleaved peptides, so that their blockades would be expected to overlap. Specifically, [F42-K51]* (1308 Å^3^) and [A1-R9]* (1260 Å^3^) had molecular volumes close to that of [T64–K73] (1369 Å^3^), whereas [G105-R112]* (1022 Å^3^) was comparable in volume to [T97-K104] (981 Å^3^). At a final concentration of 5 μM, all three peptides produced distinct blockade populations, with their respective *I/I*_*o*_ values spanning a range of 0.46 to 0.65 (Fig. 3b, Table 1). This range was comparable to that observed for the fully cleaved peptides. Notably, [G105-R112]* produced a well-defined blockade population, in contrast to its fully cleaved counterpart [G107-R112], which generated too few events for reliable characterization under the same measurement conditions. The additional N-terminal Gly-Arg sequence introduces an extra positively charged arginine residue, which may enhance electrophoretic capture of the peptide and thereby contribute to its improved detectability. The analysis of these additional reference peptides allowed further investigation of the relationship between peptide physicochemical properties and the resulting nanopore blockade characteristics.

In accordance with previously reported findings,^13,14,16^ plotting the measured *I/I*_*o*_ against peptide volume revealed a relatively linear trend in the given range, spanning residual currents from 0.48 to 0.75 (Fig. 3c). Consistent with its largest calculated volume (1369 Å^3^), peptide [T64–K73] produced the deepest blockade (*I/I*_*o*_ peak 1 = 0.476 ± 0.007, *I/I*_*o*_ peak 2 = 0.516 ± 0.006), whereas the smaller peptides [H24-R29] (878 Å^3^, *I/I*_*o*_ = 0.748 ± 0.003) and [N91-R96] (846 Å^3^, *I/I*_*o*_ = 0.741 ± 0.006) generated shallower blockades. Notably, [N91–R96] produced a marginally deeper blockade than [H24–R29], despite its smaller calculated molecular volume, suggesting that additional peptide-specific properties besides molecular volume may influence the resulting relative residual current. However, the difference between their mean *I/I*_*o*_ values (Δ*I/I*_*o*_ = 0.007) is smaller than the full width at half maximum (FWHM) of either fitted distribution (0.0114 for [H24-R29] and 0.0089 for [N91-R96]; Table 1), corresponding to only 0.6 times the FWHM of the broader distribution. For comparison, the smallest peak separation resolved as two distinct populations in this study was 0.019 for [T97-K104], corresponding to 2.3 times the FWHM. Thus, the separation between [H24-R29] and [N91-R96] is substantially smaller than the smallest separation experimentally resolved under the present conditions. As illustrated by the combined representation of the individual *I/I*_*o*_ distributions (Fig. 3e), the broader distribution of [H24–R29] encompasses that of [N91–R96], resulting in substantial overlap. Consequently, although the two peptides exhibit slightly different mean blockade amplitudes, they cannot be reliably resolved as distinct populations based on *I/I*_*o*_ alone.

In addition to the overlap between [H24-R29] and [N91-R96], the selected missed cleavage peptides ([F42-K51]*, [A1-R9]*, and [G105-R112]*) also exhibited overlapping blockade populations with fully cleaved peptides (Fig. 3d), consistent with their comparable molecular volumes. For instance, peak 1 of [F42-K51]* overlapped with peak 2 of [T64-K73], whereas their respective other blockade populations remained distinguishable. Similarly, the distribution of [G105-R112]* partially overlapped with peak 2 of [T97-K104], due to the broad distribution of the missed cleavage peptide. In addition, [A1-R9]* exhibited a particularly broad blockade distribution and a high event frequency, potentially masking blockade populations of other peptides, such as [T64-K73]. These observations illustrate the potential limitations of distinguishing individual peptides based solely on the current blockade in complex peptide mixtures.

In addition to current blockade, event frequency represents another important parameter for characterizing peptide-specific nanopore signals. Event frequencies differed by more than two orders of magnitude between synthetic peptides measured at identical concentrations (Fig. 3f). The two peptides carrying the highest net charge showed the highest event frequencies: [A1-R9]* (net charge at pH 7.5: +2.76) exhibited the highest frequency, followed by [G105-R112]* (+1.76). All remaining peptides, with net charges between +0.76 and +0.86, remained below 0.8 events·s^−1^ (Table 1). Although [T64-K73] exhibited a slightly lower event frequency than expected based on its net charge (+0.85), the overall trend is consistent with an electrophoretic contribution to peptide capture at the applied potential of -60 mV. This association is notable given the high ionic strength of the 4 M KCl electrolyte, in which long-range electrostatic interactions are expected to be strongly screened. Nevertheless, additional peptide-specific properties may also influence capture efficiency, and a contribution of electroosmotic flow (EOF) cannot be excluded.

Besides differences in blockade amplitude and event frequency between individual peptides, several peptides exhibited multiple distinct blockade populations. Contrary to the expectation of a single, sequence-specific signal per analyte, four of the synthetic peptides ([A1-R9]*, [F42-K51]*, [T64-K73], and [T97-K104]) each produced two distinct blockade populations (Fig. 3). Within each peptide, the ratio of the two peak areas was stable across replicates (RSD < 5 %, n = 3), indicating that the bimodality is a reproducible feature of the individual peptide measurements rather than a consequence of random measurement variability. Two explanations may account for two current blockade populations arising from a single molecular species: i) proline-induced conformational effects,^13,21,46^ or by ii) orientation-dependent obstruction of the conductive pore volume^8^.

1. All four peptides exhibiting two blockade populations contain at least one internal proline residue, whereas the single-population peptides either lack an internal Xaa–Pro bond ([P122-R129], whose proline occupies the N-terminus) or are among the shortest investigated ([H24-R29] and [N91-R96]). Xaa-Pro peptide bonds can adopt both *trans* and *cis* configurations (Supplementary Fig. 17), with the *cis* fraction depending on the amino acid preceding proline.^47^ NMR studies have generally reported *cis* populations of up to approximately 10% for Xaa-Pro peptide bonds, with higher values of approximately 15% for Tyr-Pro and 20% for Trp-Pro bonds.^46^ Such conformational differences may alter the effective steric volume and capture properties of a peptide, potentially giving rise to distinct blockade amplitudes. The measured peak ratios, however, are not consistent with the solution equilibrium alone. The minor population ranged from 7 % for [A1-R9]* to 34 % for [T97-K104] (Fig. 3g) and thus exceeded the reported *cis* fractions for the corresponding Xaa–Pro bonds in three of the four peptides. Because the nanopore reports an event-weighted rather than a population-weighted ratio, these values are expected to reflect conformer-dependent capture probabilities and peptide-specific interactions with the pore in addition to the underlying conformational equilibrium.
2. Peptides are structurally asymmetric along their sequence, such that entry into the pore from the N-or C-terminal end may result in different interactions with the pore lumen and consequently different current blockades. Iesu et al.^21^ reported three event populations for proline-containing peptides measured with an aerolysin nanopore under comparable electrolyte conditions, attributing two to the *cis* and *trans* conformers captured from the terminus adjacent to the proline residue and a third to entry from the opposite terminus. All peptides examined here produced exactly two populations rather than three, which may indicate that one entry orientation predominates under the present conditions, or that populations arising from the alternative orientation remain unresolved. The present dataset does not allow the contributions of peptide conformation and entry orientation to be disentangled. A more detailed investigation of the underlying mechanisms will be addressed in a separate study.

### Assignment and validation of the MBP fingerprint using the reference peptides

The established reference library of synthetic MBP peptides was used to assign the individual blockade populations observed in the SP3-processed MBP digest. The characteristic *I/I*_*o*_ values of the synthetic peptides closely matched those of the corresponding populations in the MBP fingerprint. Based on this comparison, population 1 was assigned to [T64-K73]. In agreement with the measurements of the synthetic peptide, which also yielded a comparatively low number of blockade events, this population was detected at a relatively low event frequency in the digest. Populations 2 and 3, which formed a resolved doublet in the digest, were assigned to [T97-K104], consistent with the bimodal blockade distribution observed for the synthetic reference peptide. Population 4 was attributed to [P122-R129], whereas population 5 was assigned to [H24-R29] based on its *I/I*_*o*_ value and comparatively high event frequency. Although [N91-R96] exhibited a similar *I/I*_*o*_ value, its event frequency was substantially lower, making [H24-R29] the more likely predominant contributor to population 5. To further validate these assignments, the synthetic MBP peptides were sequentially spiked into the SP3-processed MBP digest during nanopore recording (Fig. 4, Supplementary Fig. 18). Following each spike-in, the resulting blockade populations were consistent with those observed in the unspiked digest, with a maximum relative deviation of 0.32% for population 3 ([T97–K104]). The spike-in of [N91-R96] resulted in only a minor shift of population 5 towards a lower *I/I*_*o*_ value (0.005; Supplementary Fig. 18). Given the previously described overlap between the blockade distributions of [N91-R96] and [H24-R29], a contribution of [N91-R96] to population 5 can neither be quantified nor excluded, consistent with its independent identification in the MBP digest by LC-MS/MS. Taken together, each blockade population observed in the SP3-processed MBP digest could be assigned to at least one synthetic reference peptide, while LC-MS/MS identified additional MBP-derived peptides for which no distinct event population was observed in the nanopore measurements.

**Fig. 4:**
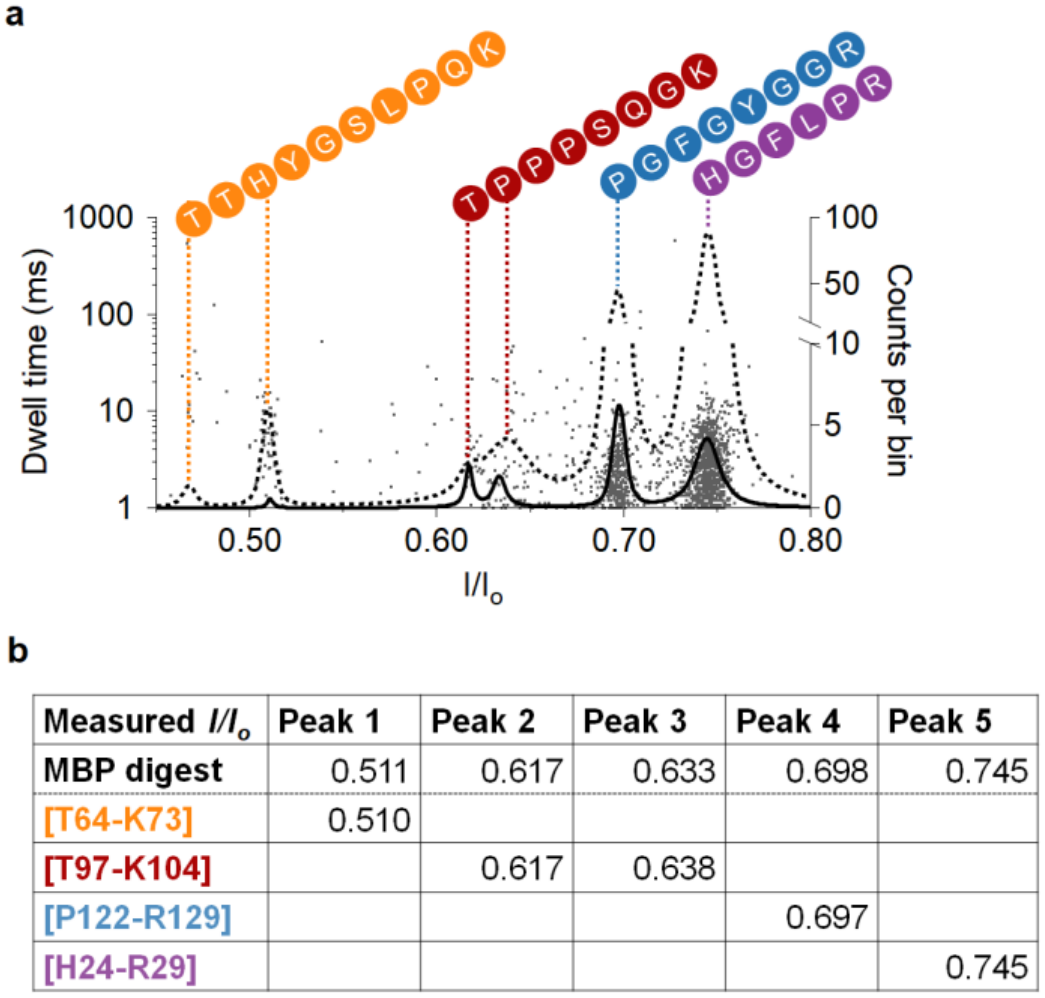
Nanopore sensing of the SP3-processed MBP digest with spike-in of synthetic MBP peptides for peptide assignment. **a** Scatter plots (grey) of the dwell time versus the relative residual current (*I/I*_*o*_) of the MBP digest (nominal final concentration: 1 µM) and synthetic MBP peptides (nominal final concentration: 5 µM) measured with AeL-R220S at -60 mV. Voigt functions^43^ (black) were fitted to histograms of the corresponding *I/I*_*o*_ distributions; the histograms are not shown for clarity. Solid lines represent the corresponding peak distributions in the unspiked MBP digest, whereas dashed lines show the distributions after sequential addition of the respective synthetic reference peptides. **b** Measured *I/I*_*o*_ values of the MBP digest compared with those of the corresponding synthetic MBP peptides.

Notably, none of the three analyzed missed cleavage peptides could be identified in the MBP fingerprint, despite producing distinct blockade populations when measured individually at a concentration of 5 µM. This may be explained by their potentially lower abundance in the digest. Based on the initial protein input, the overall peptide concentration in the digest was estimated to be approximately 1 µM (see Methods). However, this value represents an average concentration and does not account for differences in the yields of individual digestion products. In particular, missed cleavage products are not necessarily generated at the same abundance as fully cleaved peptides and may therefore be present at substantially lower concentrations. Consistent with this, the three missed cleavage peptides exhibited substantially lower LC-MS/MS signal intensities than [T97-K104], the lowest-intensity peptide detected by nanopore sensing under the applied conditions (Fig. 2d, Supplementary Table 1). Although these differences in MS signal intensity may indicate lower abundances of the missed cleavage peptides in the digest, MS intensities cannot be directly interpreted as peptide concentrations because they are also influenced by peptide-specific ionization and detection efficiencies.

The three analyzed peptides represented only a subset of the missed cleavage products identified by LC-MS/MS. Four additional peptides ([G53-R63]*, [D159-R168]*, [I152-R158]* and [L155-R161]) fell within comparable molecular volume ranges and would therefore also be expected to produce potentially overlapping blockade populations with those fully cleaved peptide. Furthermore, several other missed cleavage peptides exhibited larger molecular volumes than [T64-K73] (1369 Å^3^), the peptide with the largest molecular volume assigned to the MBP fingerprint. Based on the observed relationship between molecular volume and relative residual current (Fig. 3f), these peptides would be expected to produce deeper current blockades (*I/I*_*o*_ below 0.45). However, no corresponding events were observed in the MBP digest (Fig. 2b). Together with their generally lower MS/MS signal intensities, this suggests that these peptides may be present at insufficient abundances to generate detectable and resolvable events under the applied nanopore measurement conditions. More generally, reliable nanopore detection may require micromolar analyte concentrations, depending on the peptide and measurement conditions. Consequently, the potentially lower abundance of missed cleavage peptides may result in insufficient capture events for their detection in the digest, even though distinct blockade populations can be obtained when the corresponding synthetic peptides are measured individually at higher concentrations.

## Conclusion

In summary, this study establishes a reference-based strategy for assigning nanopore blockade populations to defined peptide sequences in a protein digest, and quantifies the limits of that assignment. Using an engineered aerolysin variant (AeL-R220S), tryptic digests of myelin basic protein (MBP) produced reproducible single-molecule peptide fingerprints, with reproducible peak positions reproducible within RSD ≤ 0.59 % across independent measurements (n =3). Direct nanopore measurement of the digest was enabled by a single-pot, solid-phase-enhanced (SP3) sample preparation, which removes reagents that destabilize lipid bilayers or interfere with Ag/AgCl electrodes,^38,39^ without additional purification steps or modifications of the experimental setup.

Individual blockade populations were assigned to peptide sequences using synthetic reference peptides established based on *in silico* digestion, physicochemical selection criteria and orthogonal LC-MS/MS identification. Comparison with the synthetic reference peptides, together with sequential spike-in experiments, confirmed the assignments of the predominant contributors to the observed blockade populations. However, peptides with similar physicochemical properties and overlapping blockade distributions, such as [H24-R29] and [N91-R96], could not be distinguished individually. Thus, assignments identify the predominant contributors to individual populations but do not exclude contributions from peptides with overlapping residual current distributions. Detectability in the pore was governed by more than abundance alone. Two peptides ([S162-R168] and [G107-R112]) gave no measurable signal despite their confirmed presence in the digest and despite meeting the applied physicochemical selection criteria, and several missed cleavage peptides produced no distinguishable population in the fingerprint although they generated characteristic blockades when measured individually at higher concentration. Event frequencies of the synthetic reference peptides spanned more than two orders of magnitude at identical concentration, with the highest values observed for the most positively charged peptides. Nanopore detection therefore depends on both the abundance of a peptide and its capture efficiency, whereas LC-MS/MS detection is governed by ionization efficiency. The two techniques consequently sample overlapping but non-congruent subsets of a digest and provide complementary rather than redundant information.

In addition, some of the analyzed MBP peptides exhibited unexpected bimodal blockade distributions. All peptides showing this behavior contained at least one proline residue, suggesting a possible contribution of proline-dependent conformational differences. However, the observations also indicate that proline alone does not account for the bimodal behavior, and orientation-dependent peptide capture may contribute to the distinct blockade populations. A more detailed investigation of the underlying mechanisms will be addressed in an independent study.

Overall, this work advances nanopore-based protein analysis beyond pattern-based fingerprinting^32,33^ by establishing a reference-based approach for peptide identification, supported by orthogonal LC-MS/MS validation and facilitated by SP3 sample preparation for direct nanopore sensing.

The present study establishes the feasibility of this workflow, prioritizing sample preparation compatibility and reference-based assignment over analytical sensitivity. Extending this approach to more complex samples will require reference databases covering a broader range of proteins and their digestion products. Furthermore, the resolution limitations identified here highlight the need for further pore engineering to improve the discrimination of peptides with overlapping residual current distributions. Ultimately, translating this approach to clinical applications will require improved analytical sensitivity, reduced sensing volumes, and workflow automation. In particular, target enrichment strategies, such as immunocapture, will need to be evaluated for their compatibility with nanopore sensing and complex biological matrices. Demonstrating matrix tolerance and reliable peptide identification in multi-protein samples will be essential next steps towards applying nanopore-based proteomics to biomarker discovery and clinical diagnostics.

## Materials and Methods

### Chemicals

Chemicals used for nanopore experiments and sample preparation were obtained from the following suppliers: Ammonium bicarbonate (ABC), chloracetamide (CAA), dithiothreitol (DTT), guanidine hydrochloride (GdmHCl), tris(2-carboxyethyl)phosphine (TCEP), Trizma™ Base and octane were purchased from Sigma-Aldrich. Acetonitrile LC-MS grade (Rotisolv®), ethanol LC-MS grade, ethylenediaminetetraacetic acid (EDTA), 4-(2-hydroxyethyl)-1-piperazineethanesulfonic acid (HEPES), hydrochloric acid (HCl), potassium chloride (KCl), sodium dodecyl sulfate (SDS), tris(hydroxymethyl)aminomethane hydrochloride (Tris-HCl) and silver chloride were purchased from Carl-Roth. Phosphate buffered saline (PBS), pH 7.2 was purchased from Gibco™. Diphytanoyl-*sn*-phosphocholine (DPhPC) was purchased from Avanti Polar Lipids as chloroform solution, aliquoted, dried to film under argon stream for long-term storage at -20 °C. Films were resolubilized in octane at 5 mg/mL before use. Trypsin, sequencing-grade was obtained from Promega as lyophilized powder, resolubilized in 1 mM HCl, aliquoted and flash frozen for long-term storage.

Chemicals used for LC-MS/MS analysis were obtained from the following suppliers: Acetonitrile (ACN, LC-MS Chromasolv™) was purchased from Honeywell, Riedel-de Haën, acetic acid (HAc, HPLC grade) from Fisher Chemical, heptafluorobutyric acid (HFBA) from Sigma-Aldrich, formic acid (FA, Optima LC-MS) from Fisher Scientific and trifluoroacetic acid (TFA, LC-MS grade) from Thermo Scientific.

### MBP *in silico* digestion and peptide calculations

The *in silico* digestion of MBP was performed by PeptideCutter (ExPASy)^50^. Expected digestion products were defined based on the predicted tryptic cleavage probabilities assigned to individual cleavage sites according to the sequence-dependent statistical model of Keil and coworker^51^ implemented in PeptideCutter. Peptide volumes were calculated based on molecular weight using the online tool Peptide Property Calculator.^44^ The theoretical net charge at pH 7.5 was calculated using the Henderson-Hasselbalch equation implemented in the MolDraw Peptide Calculator,^52^ using the pKa values reported by Thurlkill et al.^49^. The ionization states of the N-and C-termini and ionizable amino acid side chains were taken into account.

### Single-Pot, Solid-Phase enhanced Sample Preparation (SP3)

Sera-Mag™ Carboxylate-Modified Magnetic Beads [E3] and [E7] (Cytiva, Marlborough, MA) were used for SP3 digestion. Beads were equilibrated at room temperature for 30 min prior to use. Equal volumes of E3 and E7 bead suspensions (5 µL each; stock concentration: 50 mg/mL) were combined, corresponding to a total of 0.5 mg beads. The storage solution was removed using a magnetic rack and then beads were washed twice with 1 mL ddH_2_O before resuspension in 20 µL ddH_2_O to obtain a final bead concentration of 25 mg/mL.

Lyophilized myelin basic protein from bovine (MBP; Sigma-Aldrich, M1891; Uniprot: P02687) was dissolved in ddH_2_O to 10 µg/µL. 40 µg of MBP was diluted in 100 mM Tris-HCl, pH 7.5 to a final volume of 45 µL, followed by the addition of 10% SDS (f.c. 1 %). Protein denaturation was carried out at 90 °C for 3 min. Reduction and alkylation were performed by addition of 500 mM TCEP (f.c. 10 mM) and 400 mM CAA (f.c. 40 mM) and increasing of the volume to 60 µL with 50 mM ABC buffer, pH 8.5. The sample was incubated at 45 °C for 10 min.

The sample was added to the prepared bead suspension (20 µL; 25 mg/mL). Acetonitrile was added to a final concentration of 50 % (V/V) to induce protein binding. After incubation for 10 min at room temperature, the supernatant was removed. The remaining proteins bound to the beads were washed twice with 200 µL 70 % ethanol and once with 180 µL 100 % acetonitrile, followed by air drying. For enzymatic digestion, the sample was resuspended in 50 µL 50 mM ABC and trypsin was added to an enzyme-to-protein ratio of 1:50 (w/w) and incubated for 16 h at 37°C. The peptide-containing supernatant was collected and trypsin activity was terminated by heat inactivation at 90°C for 3 min.

### Desalting of samples for LC-MS/MS analysis

StageTips were assembled as described previously^53^ using two discs of C_18_ material (2215-C18, 3M) per tip. StageTips were successively conditioned with 80 μL of 100 % methanol, washed with 80 μL of 0.5 % (v/v) HAc in 80 % (v/v) ACN and equilibrated with 80 μL 0.5 % (v/v) HAc / 0.02 % (v/v) HFBA, each by centrifugation at 450 *x g* for 2-4 min. Peptides were loaded in a volume of 80 µL 0.5 % (v/v) HAc / 0.02 % (v/v) HFBA by centrifugation at 250 *x* g for 5 mins and washed once with 80 μL 0.1 % (v/v) formic acid (FA) by centrifugation at 450 *x* g for 3 - 4 min before elution of peptides in 40 μL of 0.1% (v/v) FA in 80% (v/v) ACN, using a syringe-based manual pressure device. Peptides were concentrated to a volume of 1.5 - 3.5 μL in a vacuum evaporator followed by the addition of 1.5 μL 1 % (v/v) TFA / 5 % (v/v) ACN. Finally, the total volume was adjusted to 10 μL with 0.1 % (v/v) FA of which 3.5 μL were injected for MS/MS analysis.

### LC-MS/MS measurements

LC-MS/MS analyses were performed on a QExactive Plus mass spectrometer (Thermo Fisher Scientific), coupled to an Easy nanoLC-1200 (Thermo Fisher). Chromatographic separation of peptides was performed at 32°C (temperature controlled by PRSO-V2 Sonation oven) on in-house packed fused-silica emitter microcolumns (75 μm ID, 33 cm length, Self-pack NanoLC column tube with fritted emitter, MS-WIL, ICT36007515F-50-5) filled with reverse-phase ReproSil-Pur 120 C18-AQ beads (Dr. Maisch, r119.aq.0001). Peptides were separated using a binary solvent system consisting of 0.1 % (v/v) FA (solvent A) and 80 % (v/v) ACN / 0.1 %(v/v) FA (solvent B). Peptides were loaded at 500 nL/min 0 % solvent B and then separated by the following linear gradients: from 0 % to 2 % solvent B over 5 min at 250 nL/min, from 2 % to 60 % over 40 min at 250 nL/min, from 60 % to 80 % over 4 min at 250 nL/min, and kept at 80 % for 5 min at 500 nL/min followed by an inverse gradient from 80 % to 2 % over 2 min and a re-equilibration step over 5 min at 0 % solvent B at 500 nL/min. Samples were analyzed in positive polarity DDA mode and full scan spectra were acquired at m/z 300 - 1650 with 70.000 resolution. Target values were set to 3.0e^6^ and maximum injection time of 50 ms. The 12 most intense multiply charged ions (*z* = 2 - 6) were sequentially isolated and fragmented by higher-energy collisional dissociation (HCD) with max. injection time of 60 ms and 17.000 resolution. Spray voltage was at +2.2 kV, normalized HCD collision energy (NCE) at 28 %, dynamic exclusion time at 15 s, and isolation window at 2 m/z.

### LC-MS/MS data analysis

Raw files of LC-MS/MS analyses were processed jointly using MaxQuant,^54^ version 1.6.14. For peptide identification spectra were correlated with Myelin basic protein (Uniprot identifier P02687) and a common contaminants FASTA using the Andromeda^55^ search engine embedded in MaxQuant. Cysteine carbamidomethylation was set as a fixed modification and N-terminal acetylation, methionine oxidation and deamidation at asparagine or glutamine as variable modifications with deamidation at asparagine or glutamine excluded within the “First search” option. The enzyme specificity was set to Trypsin/P (enabling cleavage before proline) and a maximum of two missed cleavages were allowed. The minimum peptide length was set to 6 amino acids. A false discovery rate of 1 % was applied to both PSM and protein lists using the decoy mode “Revert.” The option “second peptides” was disabled and the option “Match between Runs” was left unchecked.

### Nanopore recording

An engineered variant of the biological aerolysin (AeL) pore was used for all experiments. Biosynthesis of the inactive pAeL-R220S was carried out as described by Ensslen et al.^14,42^ 1 µL of pAeL-R220S (1 mg/mL) was diluted with 8 µL PBS, 100 µM DTT, 100 µM EDTA, pH 7.5. Addition of 1 µL trypsin (12.5 mg/mL, 1 mM HCl) was used to activate the proaerolysin. After mixing, the solution was incubated for 15 min at room temperature. Nanopore recordings were performed using the E16 amplifier (Elements SRL, Cesena, Italy) integrated into the Orbit16TC parallel bilayer recording device (Nanion Technologies, Munich, Germany) using MECA16 microelectrode cavity array chips (Ionera Technologies, Freiburg, Germany) with a cavity diameter of 100 µm. The temperature was set to 18 °C. Bilayer formation and pore insertion were controlled by the EDR4 software (Elements SRL, Cesena, Italy). The EDR4 software settings were set as follows: voltage command filter: 1 kHz, current range: 2 nA, sampling rate: 20 kHz. The MECA16 chip was filled with 200 µL 4 M KCl, 25 mM Tris-Base, pH 7.5, saturated with AgCl. Lipid bilayers were formed using a lipid-coated air-bubble method^56–58^ with 5 mg/mL of DPhPC in octane. The activated aerolysin solution was added to the common *cis* compartment of the chip to a final concentration of 0.35 µg/mL. Although the MECA16 enables parallel monitoring of up to 16 cavities, each measurement presented in this study was performed on single cavities, each containing a single aerolysin pore. Independent technical replicates were obtained using new chips. After successful pore insertion, 5 µL of the SP3-processed MBP digest was added to the *cis* compartment and measured at an applied potential of -60 mV for 10 minutes. The MBP digest volume and the amount introduced into the chip were calculated based on the initial weighted MBP amount. Assuming complete proteolytic digestion and no peptide loss, this corresponds to a final MBP concentration of 1 µM during nanopore measurements.

Additional measurements at potentials ranging from -40 mV up to -90 mV were used to evaluate the voltage dependency of event frequency (events·s^-1^) and dwell time (ms).

### Nanopore analysis of synthetic MBP peptides

All synthetic MBP peptides were provided by ProteoGenix (Schiltigheim, France). According to the manufacturer, the synthetic peptides had a purity of > 94 %, as determined by HPLC analysis, and their molecular weight was confirmed by MS analysis (Supplementary Table 3). Lyophilized peptides were reconstituted in 5 mM HEPES, pH 7.5, based on the peptide molecular weights to obtain 1 mM stock concentrations. 1 µL of each peptide was added to the *cis* side of the chip at a final concentration of 5 µM.

### Nanopore data analysis

Data were recorded with the Elements EDR4-Software. The data files in Axon binary format 2 were analyzed with in-house software written in C# using a sliding window baseline correction algorithm and threshold-based peak detection. Events with dwell times < 1 ms were excluded from subsequent analyses because their durations could not be reliably resolved under the applied recording conditions. Further analysis and graphing were performed using IgorPro9 (Wavemetrics, Portland, OR). Scatter plots of the dwell time versus the relative residual current (*I/I*_*o*_) of individual blockade events were generated. Histograms were constructed using a bin width of 0.001, and Voigt functions^43^ were fitted to the resulting distributions using the Multipeak Fitting procedure in Igor to determine the peak maxima. Event frequencies for individual peptides were determined by counting the blockade events associated with each Voigt-fitted peak and dividing the number of events by the total measurement duration (s), yielding the event frequency in events·s^-1^.

## Author contributions

‡These authors contributed equally.

- **Carolin Hensel‡:** Conceptualization, Methodology, Investigation: Established, conducted and analyzed nanopore measurements, adapted the SP3 workflow for nanopore analysis of MBP, Visualization, Writing (original draft)
- **Carolin Kleber‡:** Conceptualization, Methodology, Review & Editing
- **Ida Suppanz:** Investigation: Performance and analysis of LC-MS/MS experiments
- **Atefeh Akbari:** Investigation: Pilot experiments and initial identification of MBP-related nanopore signals for peptides [T64-K73], [H24-R29] and [N91-R96]
- **Sebastian Gutsch:** Development and implementation of software for nanopore data analysis
- **Tobias Hutzenlaub:** Supervision, Funding Acquisition, Review & Editing
- **Tobias Ensslen:** Supervision, Review & Editing
- **Jan C. Behrends:** Project Conceptualization, Supervision, Review & Editing, Funding Acquisition
- **Felix von Stetten:** Project Conceptualization, Supervision, Review & Editing
- **Gerhard Mittler:** Supervision and conceptualization of LC-MS/MS experiments, Review & Editing
- **Jan-Niklas Klatt:** Supervision, Project Administration, Review & Editing
- **Andreas Schreiber*:** Conceptualization, Supervision, Project Administration, Review & Editing

## Acknowledgements

- **Declaration of Generative AI and AI-assisted Technologies in the Writing process**. AI tool GPT-5.6 (OpenAI) aided manuscript editing. Authors created all original content, reviewed and revised AI suggestions, and assume full accountability for the final published version.
- **Funding sources**. This work was financially supported by the BMFTR within the project ProNanoPep (project number 031B1120A/B).

## Supporting information

Further information is available in the Supplementary Information.

## Code availability

The custom software used for baseline correction and event detection is available from the corresponding author on reasonable request.

## Data availability

The mass spectrometry proteomics data have been deposited to the ProteomeXchange Consortium via the PRIDE partner repository with dataset identifier PXD086193.^59^ The authors declare that all data supporting the findings of this study are available within the paper and its supplementary information files. The dataset for nanopore measurement underlying the figures and tables presented in this study are available within the article and its Supplementary Information.

