## Supplementary for "Reference-based assignment of aerolysin nanopore signals to peptide sequences from tryptic protein digests"

*‡ Equal authorship*

*[1] Laboratory for MEMS Applications, IMTEK - Department of Microsystems Engineering, University of Freiburg, Germany*

*[2] Max-Planck Institute for Immunobiology and Epigenetic, Freiburg, Germany*

*[3] Laboratory for Membrane Physiology and Technology, Department of Physiology, University of Freiburg, Germany*

*[4] Hahn-Schickard gGmbH, Freiburg, Germany*

### SI Notes:

**Note 1: Nanopore analysis of MBP in-solution digest.** Consistent with the known membrane-disrupting effects of detergents, preliminary nanopore measurements of unpurified conventional in-solution digests of proteins led to rapid membrane destabilization and loss of stable open-pore currents, highlighting the need for efficient removal of nanopore-incompatible reagents.

The in-solution digestion protocol was adapted from Promega (Rapid-Digestion Trypsin/Lys-C Kits).[1] Lyophilized myelin basic protein (MBP; Sigma-Aldrich, M1891) was dissolved in dd$H_2O$ to 0.5 µg/µL. 5 µg of MBP were used for the in-solution digestion and denatured by the addition of 30 µL denaturation buffer (50 mM Tris, 8 M guanidine hydrochloride (GuHCl), pH 8), resulting in a final GuHCl concentration of 6 M. Reduction was performed by addition of DTT (f.c. 2.5 mM), followed by incubation at 95°C for 20 min. Cysteine residues were alkylated by addition of chloracetamide (CAA; f.c. 10 mM) and incubation for 1 h. Remaining alkylation reagent was quenched by addition of DTT (f.c. 10 mM) and incubation for 1 h. The denatured protein solution was diluted with digestion buffer (50 mM Tris, 1 mM $CaCl_2$, pH 7.6) to reduce the GuHCl concentration to 0.8 M. Proteolytic digestion was done by addition of trypsin (0.1 µg, enzyme-to-substrate ratio 1:50) and incubation at 37°C for 2 h. The reaction was terminated by addition of trichloroacetic acid (TCA; f.c. 10 %). Successful digestion was confirmed by LC-MS/MS analysis, which demonstrated the presence of tryptic peptides. 5 µL of the digest were added to the *cis* compartment of the MECA16, containing 200 µL KCl, pH 7.5. However, direct application of the resulting digest to the nanopore measurement setup led to loss of stable open-pore currents, presumably owing to residual nanopore-incompatible reagents.

**Note 2: Nanopore analysis of trypsin autolysis products.** To assess whether nanopore signals could originate from trypsin autolysis products, a trypsin control was prepared, exceeding the amount used in the SP3 digestion (0.8 µg trypsin). Briefly, 5 µg of sequence-grade trypsin were incubated in 50 mM ABC (final volume, 50 µL) for 16 h at 37°C. Trypsin activity was then terminated by heat inactivation at 90°C for 3 min. The sample was directly analyzed using the aerolysin nanopore at a nominal final trypsin concentration of 0.1 µM, resulting in only rare current blockades.

### SI Tables:

**Table 1: MBP peptides identified by LC-MS/MS analysis of SP3-processed MBP digests.** Reverse-sequence entries and contaminant entries were removed. Prior to bead binding samples were denatured by heat (3 min, 90°C) either without SDS (three replicates, Rep1 - Rep3) or in the presence of 1 % SDS (four replicates, Rep1 - Rep4) using varying protein input (20 - 80 µg). The table lists the peptide sequences and corresponding intensities identified in the SDS Rep1 sample, which was used for all nanopore measurements in the main paper. Asterisks (*) denote peptides containing missed cleavages. Peptide intensities are color-coded from low (red) to high (green) intensity. Nanopore measurements of SDS Rep2 - Rep4 are illustrated in SI Fig. 4. The complete LC-MS/MS datasets, including all replicates are provided in the accompanying excel file.

| **Peptide** | **Sequence** | **Nanopore Ref.: Intensity SDS Rep1** |
|---|---|---|
| [A1-R9]* | AAQKRPSQR | 2.76E+07 |
| [S10-R23]* | SKYLASASTMDHAR | 1.55E+08 |
| [Y12-R23] | YLASASTMDHAR | 1.41E+11 |
| [H24-R29] | HGFLPR | 9.66E+10 |
| [H30-R41]* | HRDTGILDSLGR | 4.18E+09 |
| [D32-R41] | DTGILDSLGR | 8.32E+10 |
| [F42-R47] | FFGSDR | 8.15E+10 |
| [F42-R51]* | FFGSDRGAPK | 3.25E+08 |
| [G53-R63]* | GSGKDGHHAAR | 8.47E+06 |
| [T64-R73] | TTHYGSLPQK | 8.98E+10 |
| [T64-R78]* | TTHYGSLPQKAQGHR | 3.93E+06 |
| [A74-R90]* | AQGHRPQDENPVVHFFK | 1.11E+09 |
| [P79-R90] | PQDENPVVHFFK | 1.94E+10 |
| [N91-R96] | NIVTPR | 7.48E+10 |
| [N91-R104]* | NIVTPRTPPPSQGK | 1.28E+09 |
| [T97-R104] | TPPPSQGK | 2.65E+09 |
| [G105-R112]* | GRGLSLSR | 1.58E+07 |
| [G107-R112] | GLSLSR | 3.24E+10 |
| [F113-R121] | FSWGAEGQK | 4.16E+10 |
| [F113-R129]* | FSWGAEGQKPGFGYGGR | 5.83E+08 |
| [P122-R129] | PGFGYGGR | 6.09E+10 |
| [P122-R134]* | PGFGYGGRASDYK | 4.20E+07 |
| [G139-R151]* | GLKGHDAQGTLSK | 8.84E+08 |
| [G142-R151] | GHDAQGTLSK | 5.54E+10 |
| [G142-R154]* | GHDAQGTLSKIFK | 1.90E+07 |
| [I152-R158] | IFKLGGR | 4.95E+07 |
| [I152-R161]* | IFKLGGRDSR | 8.73E+06 |
| [L155-R161]* | LGGRDSR | 7.50E+07 |
| [D159-R168]* | DSRSGSPMAR | 1.05E+08 |
| [S162-R168] | SGSPMAR | 4.25E+09 |

**Table 2: Resulting peak populations observed in the MBP digest.** The normalized residual current ($I/I_o$), full width at half maximum (FWHM) and event frequency (at f.c. 1 µM) are reported as mean ± standard deviation (SD) from three independent measurements (n = 3), performed at an applied voltage of -60 mV. Measurements were recorded for approximately 600 s. For Peak 1, only two independent measurements (n = 2) were included in the calculation of the mean $I/I_o$ and event frequency due to the low number of detected events.

| Peak | $I/I_o$ | FWHM | Frequency (events·s$^{-1}$) |
|---|---|---|---|
| 1 | 0.512 ± 0.003 | 0.0048 ± 0.0017 | 0.010 ± 0.001 |
| 2 | 0.621 ± 0.004 | 0.0101 ± 0.0030 | 0.044 ± 0.010 |
| 3 | 0.638 ± 0.004 | 0.0085 ± 0.0022 | 0.040 ± 0.010 |
| 4 | 0.700 ± 0.003 | 0.0120 ± 0.0021 | 0.128 ± 0.020 |
| 5 | 0.749 ± 0.003 | 0.0172 ± 0.0042 | 0.147 ± 0.026 |

**Table 3: Synthetic MBP peptides used for nanopore reference measurements.** Peptide sequences, molecular weights and purities of the synthetic peptides obtained from ProteoGenix. Purities were determined by HPLC* and molecular weights were confirmed by mass spectrometry (MS)**, as reported by the manufacturer.

| Peptide | Sequence | Purity (%)* | Mass (g·mol$^{-1}$)** |
|---|---|---|---|
| [T64-K73] | TTHYGSLPQK | 94.17 | 1131.36 |
| [F42-K51]* | FFGSDRGAPK | 96.31 | 1081.18 |
| [A1-R9]* | AAQKRPSQR | 95.50 | 1041.17 |
| [G105-R112]* | GRGLSLSR | 97.25 | 844.96 |
| [T97-K104] | TPPPSQGK | 96.53 | 810.89 |
| [P122-R129] | PGFGYGGR | 95.36 | 809.87 |
| [H24-R29] | HGFLPR | 95.86 | 725.92 |
| [S162-R168] | SGSPMAR | 95.89 | 704.80 |
| [N91-R96] | NIVTPR | 94.00 | 698.81 |
| [G107-R112] | GLSLSR | 96.89 | 631.72 |

### SI Figures:

AAQK R PSQR SK YLASASTMDHAR HGFLPR HR DTGILDSLGR FFGSDR GAPK R GSGK DGHHAAR TTHYGSLPQK AQGH**R*****P**QDENPVVHFFK NIVTPR TPPPSQGK GR GLSLSR FSWGAEGQ**K**P**GFGYGGR ASDYK SAHK GLK GHDAQGTLSK IFK LGGR DSR SGSPMAR R

**SI Fig. 1: *In silico* tryptic digestion of myelin basic protein (MBP) for the identification of suitable reference peptides for nanopore analysis.** Trypsin cleavage sites (lysine, K, and arginine, R) are highlighted in red. Cleavage sites located in close proximity to adjacent cleavage sites, for which a lower cleavage probability is expected, are not highlighted. Furthermore, cleavage sites followed by proline (K-P or R-P) are less accessible to trypsin and therefore exhibit a reduced cleavage probability. The two exceptions are indicated by asterisks: (*) represents a cleavage probability of 24 %, whereas (**) represents a cleavage probability of 37%. The resulting peptides highlighted in the figure are predicted to be generated with high probability. Peptides highlighted in light blue do not meet the physicochemical criteria required for nanopore detection (length of 4-20 amino acids and a positive net charge). In contrast, dark blue peptides fulfill both criteria and were therefore selected as reference peptides for peak assignment (seven peptides in total). Cleavage probabilities were obtained using PeptideCutter (ExPASy).[5]

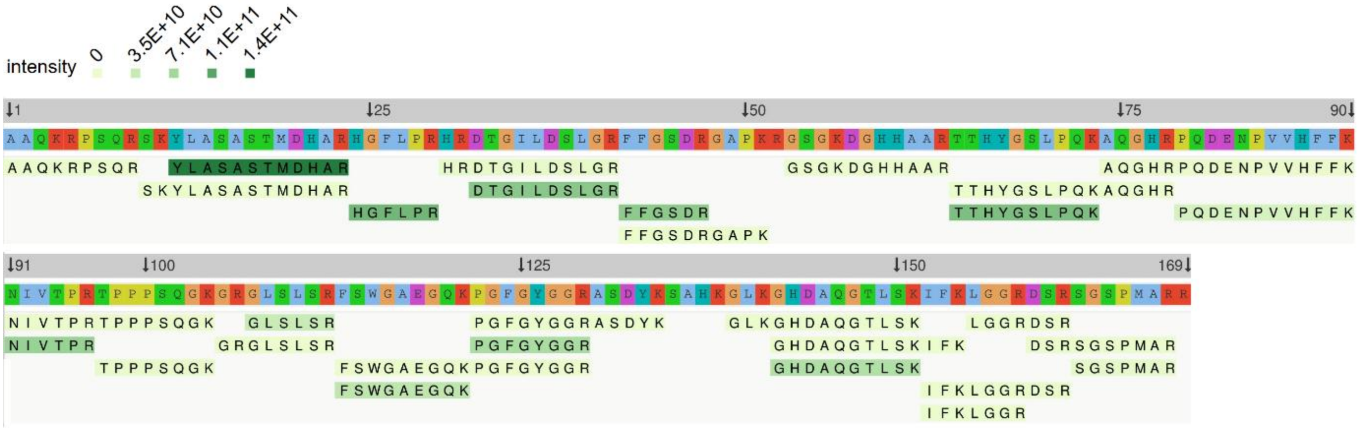


**SI Fig. 2: Peptide identification and sequence coverage obtained by LC-MS/MS analysis of the MBP digest.** The data were visualized using the Peptigram tool.[6] The protein sequence is shown with identified peptides mapped to their respective positions. Peptide intensities are represented by color intensity, with peptides containing missed cleavages resulting in a lighter shade than peptides with zero missed cleavages.

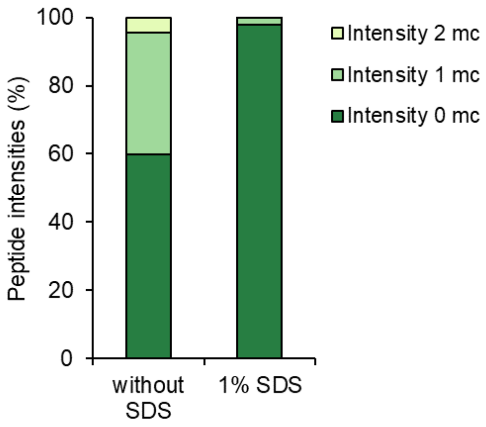


**SI Fig. 3: Missed cleavage distribution of identified peptides in LC-MS/MS analysis of samples denatured in the absence or presence of 1 % SDS (f.c.), showing the relative abundance of peptides with zero, one and two missed cleavages after digestion.**

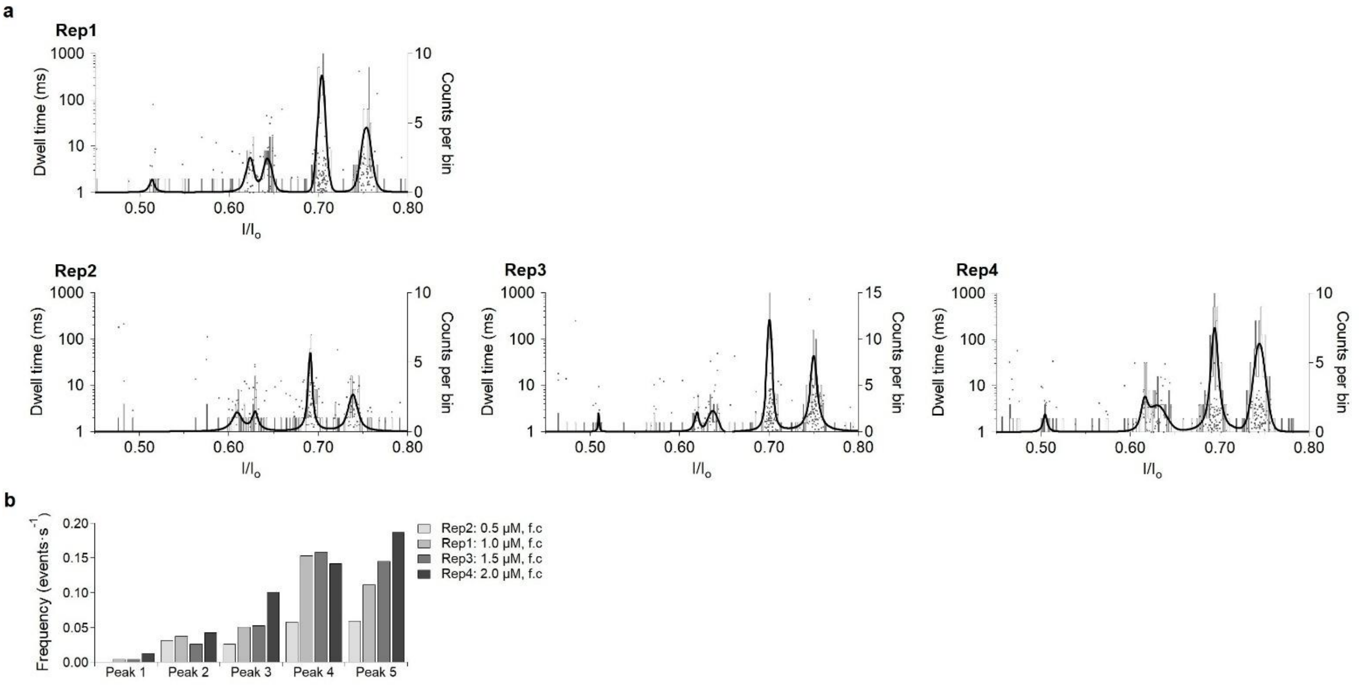


**SI Fig. 4: Nanopore sensing of the SP3-processed MBP digests with varying protein input. a** Scatter plots (grey) of the dwell time versus the relative residual current ($I/I_o$) of the SP3-processed MBP digest measured with AeL-R220S at -60 mV. Voigt functions (black) were fitted to histograms of the corresponding $I/I_o$ distributions. Four independent SP3 digests (SDS Rep1-Rep4) were prepared using increasing amounts of MBP (Rep1: 40 µg; Rep2: 20 µg; Rep3: 60 µg; Rep4: 80 µg) to identify the protein input required for reliable nanopore detection while minimizing sample consumption. SDS Rep1, prepared from 40 µg MBP, was used for nanopore measurements presented in the main paper. LC-MS/MS results for the corresponding digests are listed in Supplementary Table 1. **b** Event frequencies of the five blockade populations detected in Rep1–Rep4 at final peptide concentrations ranging from 0.5 to 2.0 µM in the nanopore measurement. Event frequencies increased for all five populations with increasing protein input and, consequently, increasing peptide concentration.

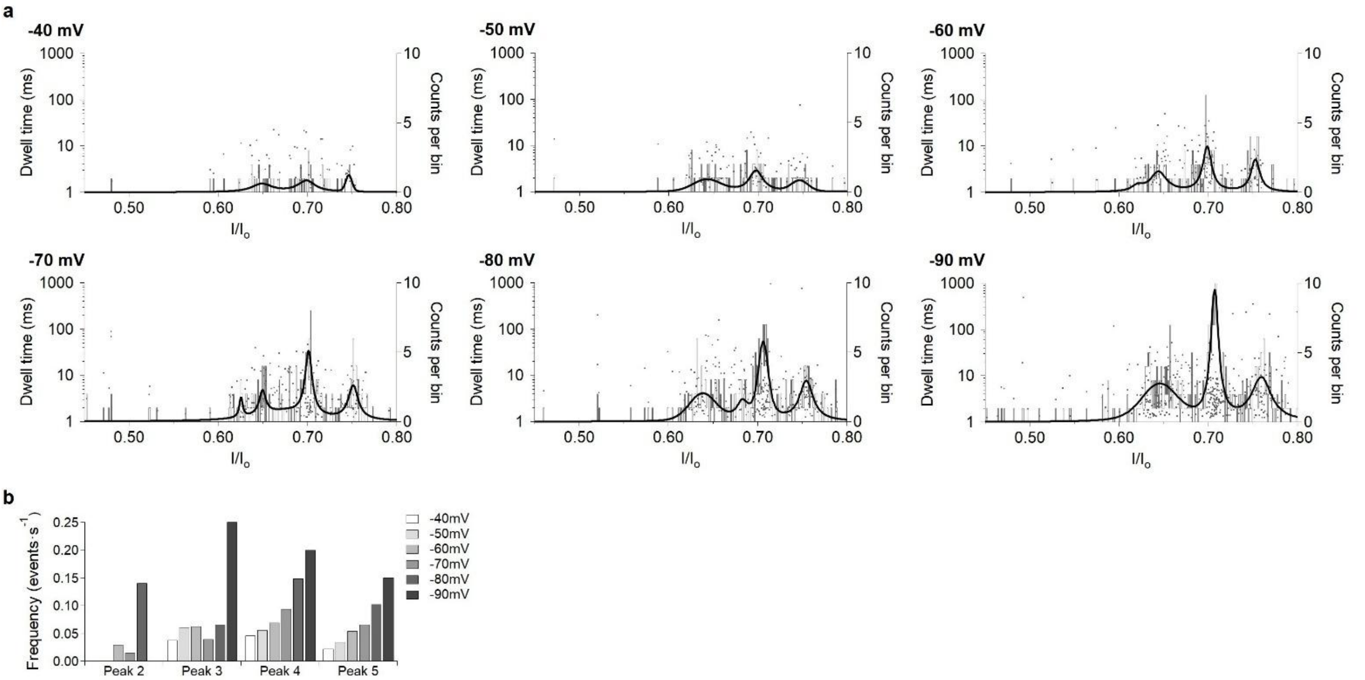


**SI Fig. 5: Optimization of nanopore sensing of the SP3-processed MBP digest for peak separation.** Nanopore analysis of the MBP digest (final concentration: 1 µM) measured with AeL-R220S at applied potentials from -40 mV to -90 mV. **a** Scatter plots (grey) of the dwell time versus the relative residual current ($I/I_o$) of the MBP digest. Voigt functions (black) were fitted to histograms of the corresponding $I/I_o$ distributions. Optimal potential for peak separation showed an applied potential of -60 mV. **b** Event frequencies of the five blockade populations detected in MBP digest at different applied potentials.

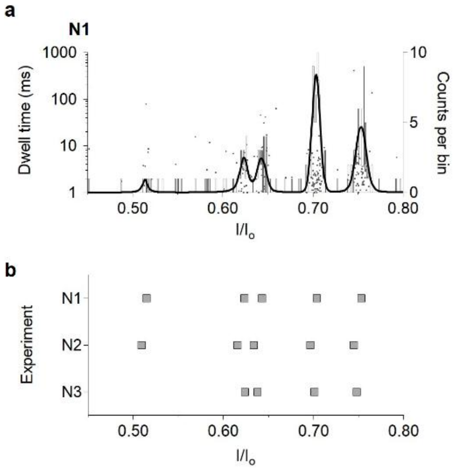


**SI Fig. 6: Nanopore sensing of the SP3-processed MBP digest. a** Scatter plots (grey) of the dwell time versus the relative residual current ($I/I_o$) of the MBP digest (final concentration: 1 µM) measured with AeL-R220S at -60 mV. Voigt functions (black) were fitted to histograms of the corresponding $I/I_o$ distributions. Analysis of the same MBP digest (Rep1, Supplementary Table 1) were performed in three independent nanopore measurements (N1 - N3). **b** Plot of the $I/I_o$ values measured in each independent measurement.

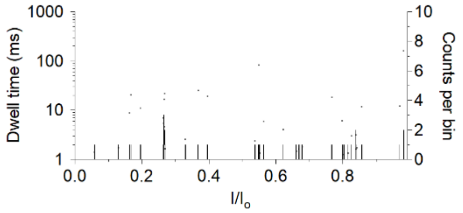


**SI Fig. 7: Nanopore sensing of trypsin autolysis products.** Scatter plots (grey) of the dwell time versus the relative residual current ($I/I_o$) of the resulting peptides (final concentration: 0.1 µM) measured with AeL-R220S at -60 mV. Peak fitting to histograms (black) of the corresponding $I/I_o$ distributions was not possible due to the low number of detected events.

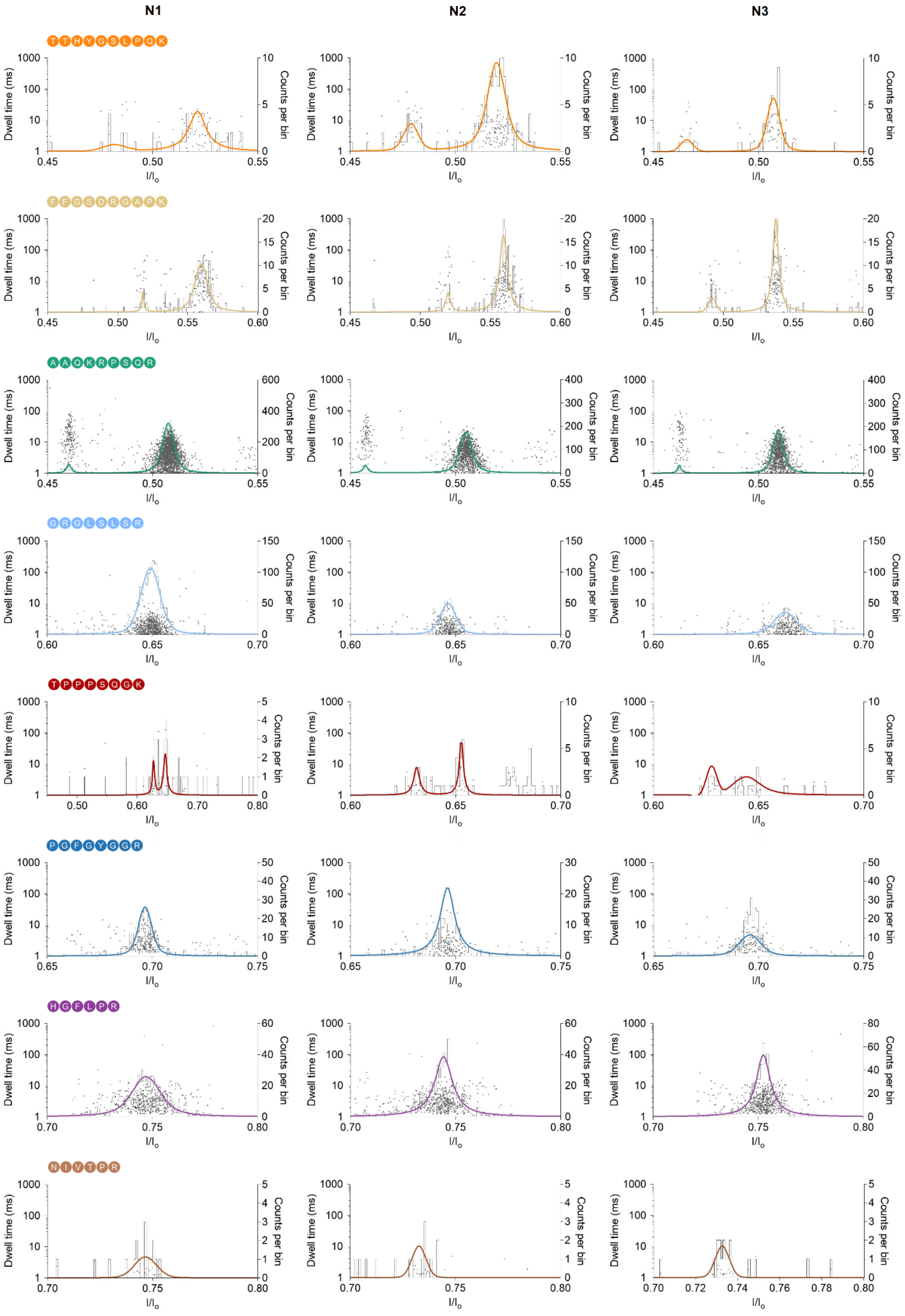
N1
N2
N3
TTHYGSLPQK
FFGSDRGAPK
AAQKRPSQR
GRGLSLSR
TPPPSQGK
PGFGYGGR
HGFLPR
NIVTPR
Dwell time (ms)
Counts per bin
I/Io

**SI Fig. 8: Nanopore sensing of synthetic MBP peptides.** Scatter plots (grey) of the dwell time versus the relative residual current ($I/I_o$) of synthetic MBP peptides (final concentration: 5 µM) measured with AeL-R220S at -60 mV. Voigt functions (colored) were fitted to histograms (black) of the corresponding $I/I_o$ distributions. Measurements were performed in triplicate (N1 - N3).

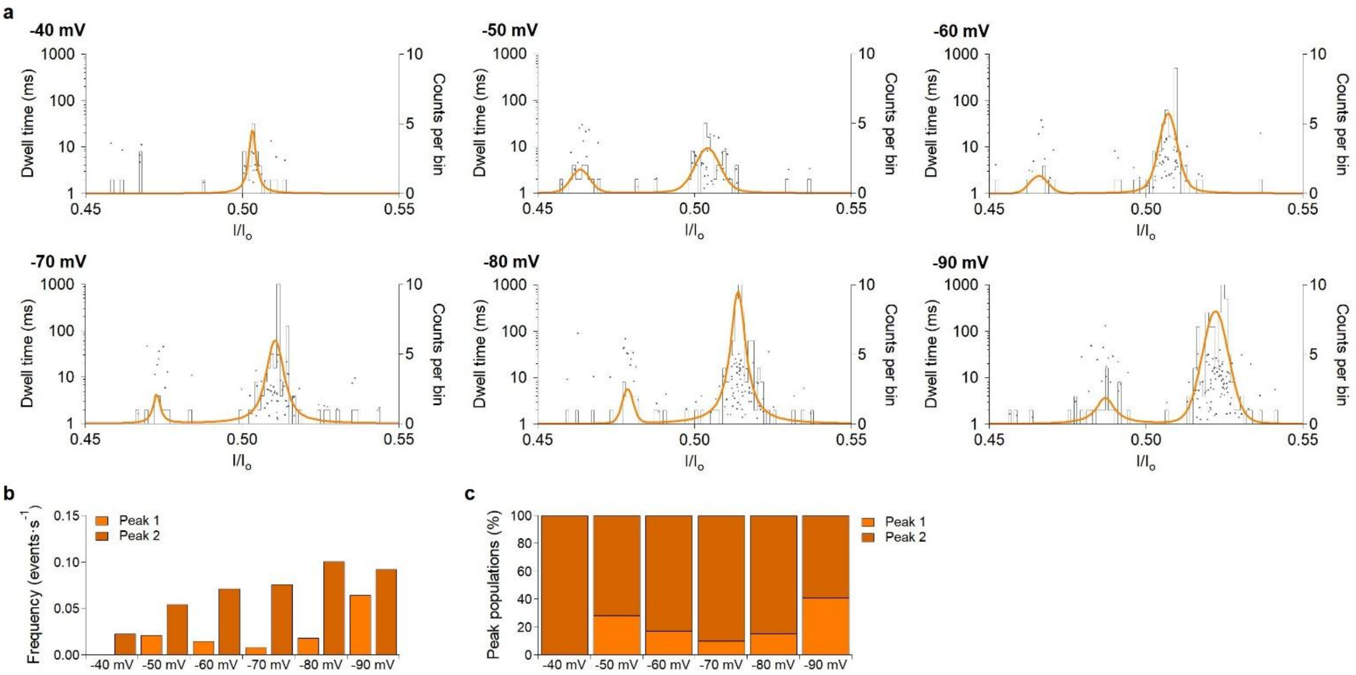


**SI Fig. 9: Nanopore sensing of synthetic MBP peptide [T64-K73].** Analysis of the peptide (final concentration: 5 µM) measured with AeL-R220S at applied potentials ranging from -40 mV to -90 mV. **a** Scatter plots (grey) of the dwell time versus the relative residual current ($I/I_o$). Voigt functions (colored) were fitted to histograms (black) of the corresponding $I/I_o$ distributions. **b** Event frequencies of the blockade populations detected at different applied potentials. **c** Relative contributions of the two peak populations forming the doublet at different applied potentials.

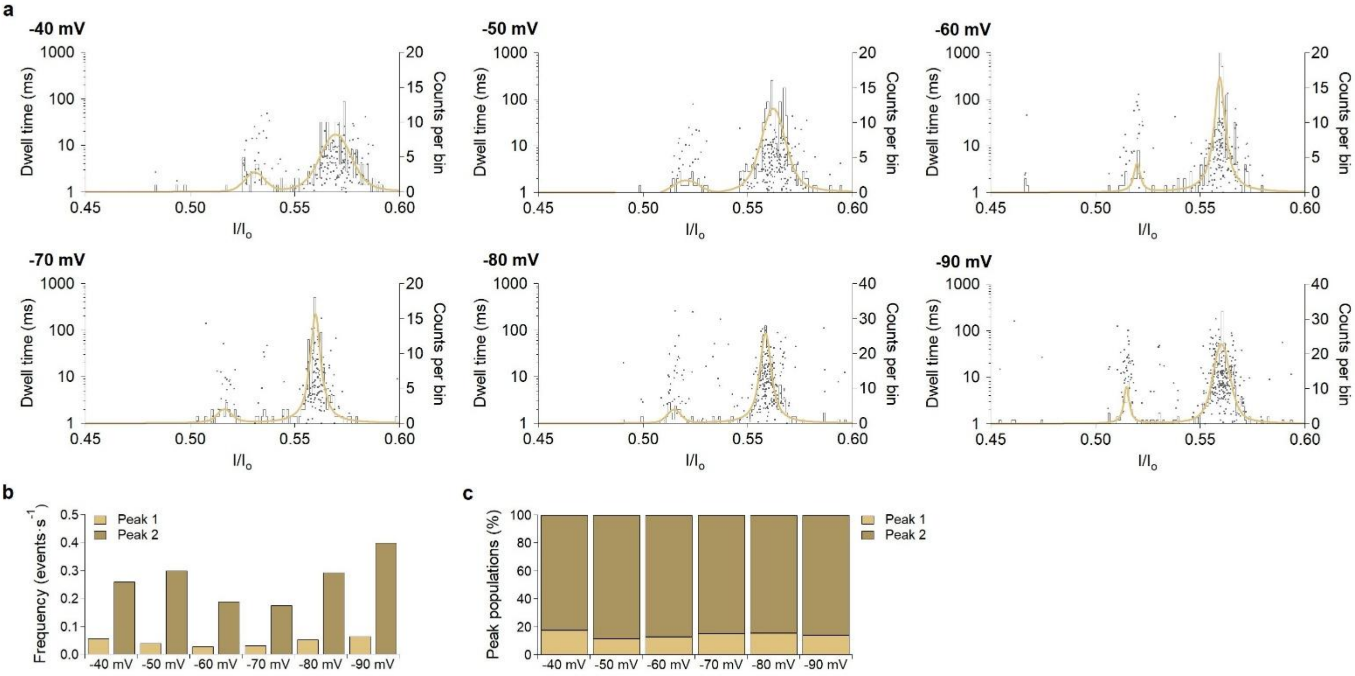


**SI Fig. 10: Nanopore sensing of synthetic MBP peptide [F42-K51]*.** Analysis of the peptide (final concentration: 5 µM) measured with AeL-R220S at applied potentials ranging from -40 mV to -90 mV. **a** Scatter plots (grey) of the dwell time versus the relative residual current ($I/I_o$). Voigt functions (colored) were fitted to histograms (black) of the corresponding $I/I_o$ distributions. **b** Event frequencies of the blockade populations detected at different applied potentials. **c** Relative contributions of the two peak populations forming the doublet at different applied potentials.

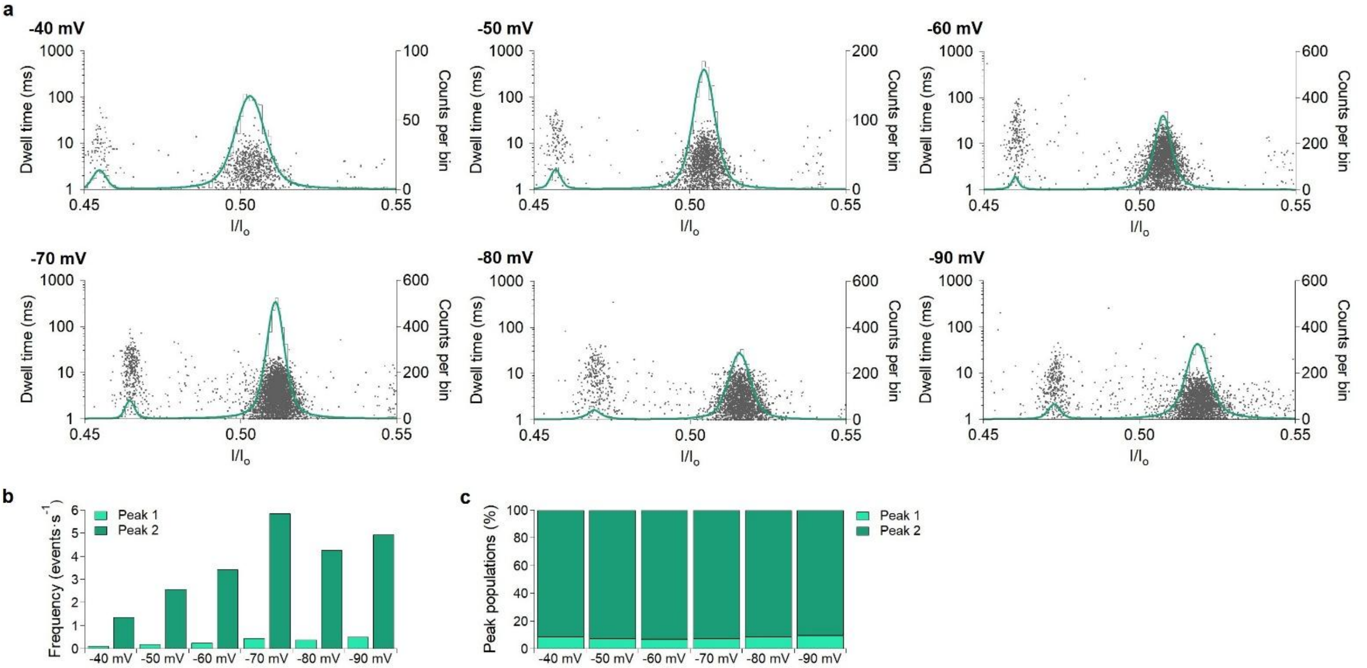


**SI Fig. 11: Nanopore sensing of synthetic MBP peptide [A1-R9]*.** Analysis of the peptide (final concentration: 5 µM) measured with AeL-R220S at applied potentials ranging from -40 mV to -90 mV. **a** Scatter plots (grey) of the dwell time versus the relative residual current ($I/I_o$). Voigt functions (colored) were fitted to histograms (black) of the corresponding $I/I_o$ distributions. **b** Event frequencies of the blockade populations detected at different applied potentials. **c** Relative contributions of the two peak populations forming the doublet at different applied potentials.

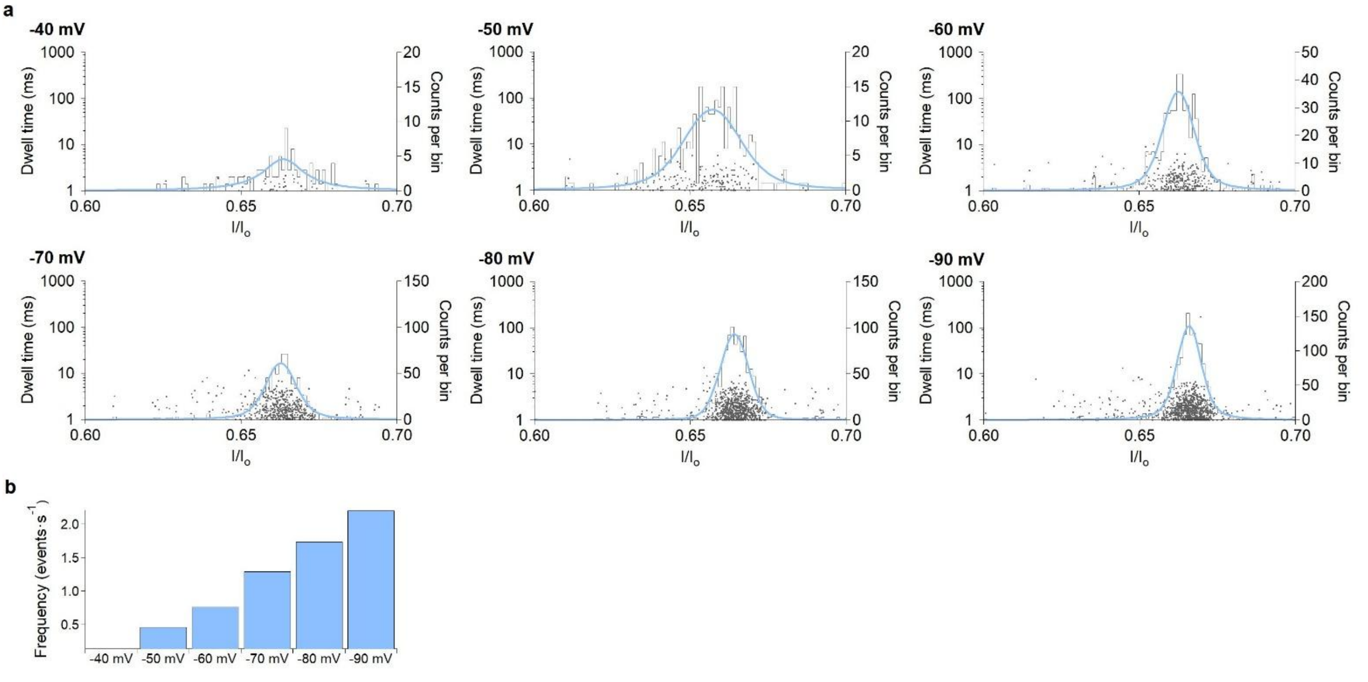


**SI Fig. 12: Nanopore sensing of synthetic MBP peptide [G105-R112]*.** Analysis of the peptide (final concentration: 5 µM) measured with AeL-R220S at applied potentials ranging from -40 mV to -90 mV. **a** Scatter plots (grey) of the dwell time versus the relative residual current ($I/I_o$). Voigt functions (colored) were fitted to histograms (black) of the corresponding $I/I_o$ distributions. **b** Event frequencies of the blockade populations detected at different applied potentials.

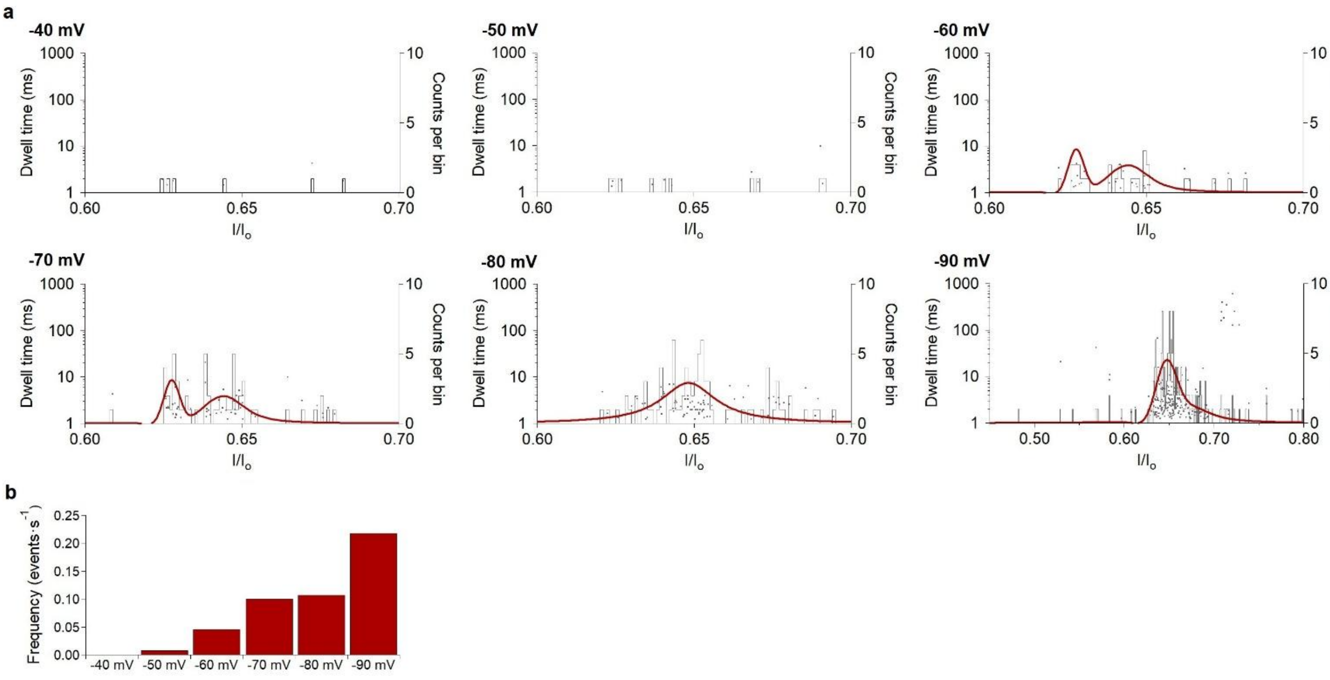


**SI Fig. 13: Nanopore sensing of synthetic MBP peptide [T97-K104].** Analysis of the peptide (final concentration: 5 µM) measured with AeL-R220S at applied potentials ranging from -40 mV to -90 mV. **a** Scatter plots (grey) of the dwell time versus the relative residual current (*I/Io*). Voigt functions (colored) were fitted to histograms (black) of the corresponding *I/Io* distributions. No peak fitting was performed at -40 mV due to the low number of detected events. At intermediate applied potentials (-50 to -70 mV), two distinct peaks were resolved. At higher applied potentials (-80 and -90 mV), the peaks could no longer be resolved, resulting in a single broadened distribution. **b** Event frequencies of the blockade populations detected at different applied potentials.

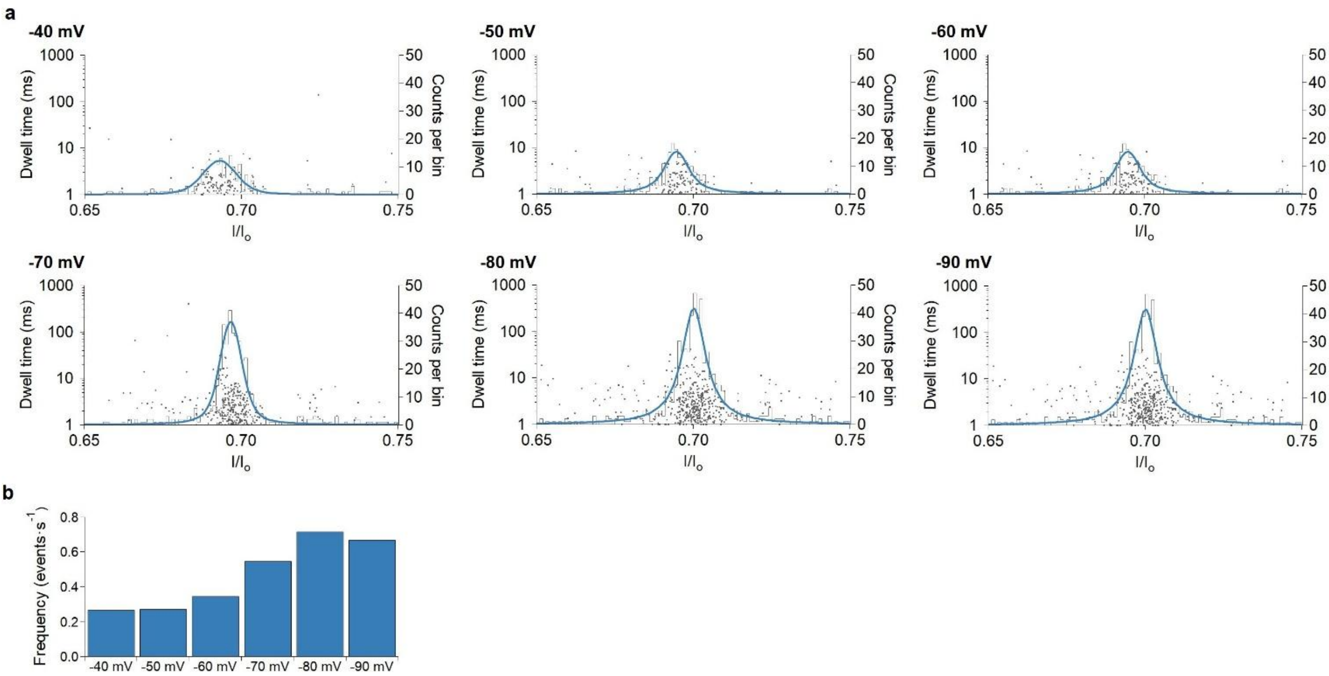


**SI Fig. 14: Nanopore sensing of synthetic MBP peptide [P122-R129].** Analysis of the peptide (final concentration: 5 µM) measured with AeL-R220S at applied potentials ranging from -40 mV to -90 mV. **a** Scatter plots (grey) of the dwell time versus the relative residual current (*I/Io*). Voigt functions (colored) were fitted to histograms (black) of the corresponding *I/Io* distributions. **b** Event frequencies of the blockade populations detected at different applied potentials.

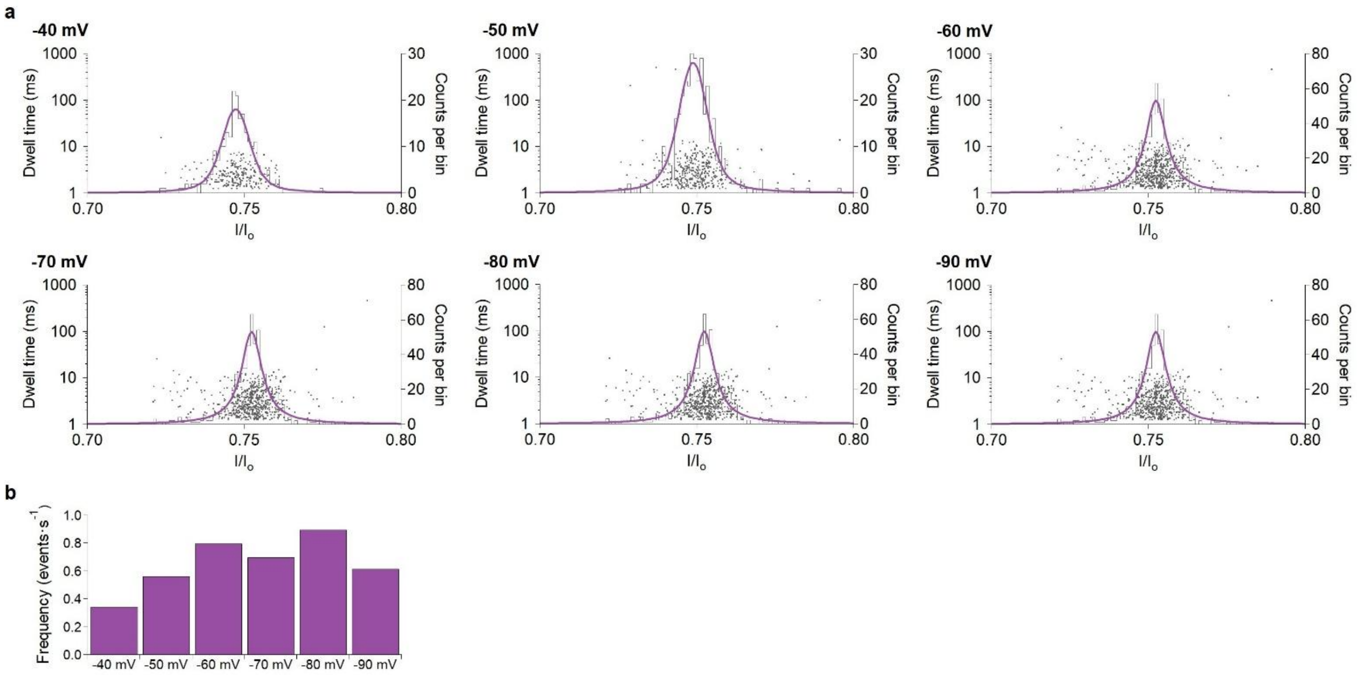


**SI Fig. 15: Nanopore sensing of synthetic MBP peptide [H24-R29].** Analysis of the peptide (final concentration: 5 µM) measured with AeL-R220S at applied potentials ranging from -40 mV to -90 mV. **a** Scatter plots (grey) of the dwell time versus the relative residual current (*I/I*$_o$). Voigt functions (colored) were fitted to histograms (black) of the corresponding *I/I*$_o$ distributions. **b** Event frequencies of the blockade populations detected at different applied potentials.

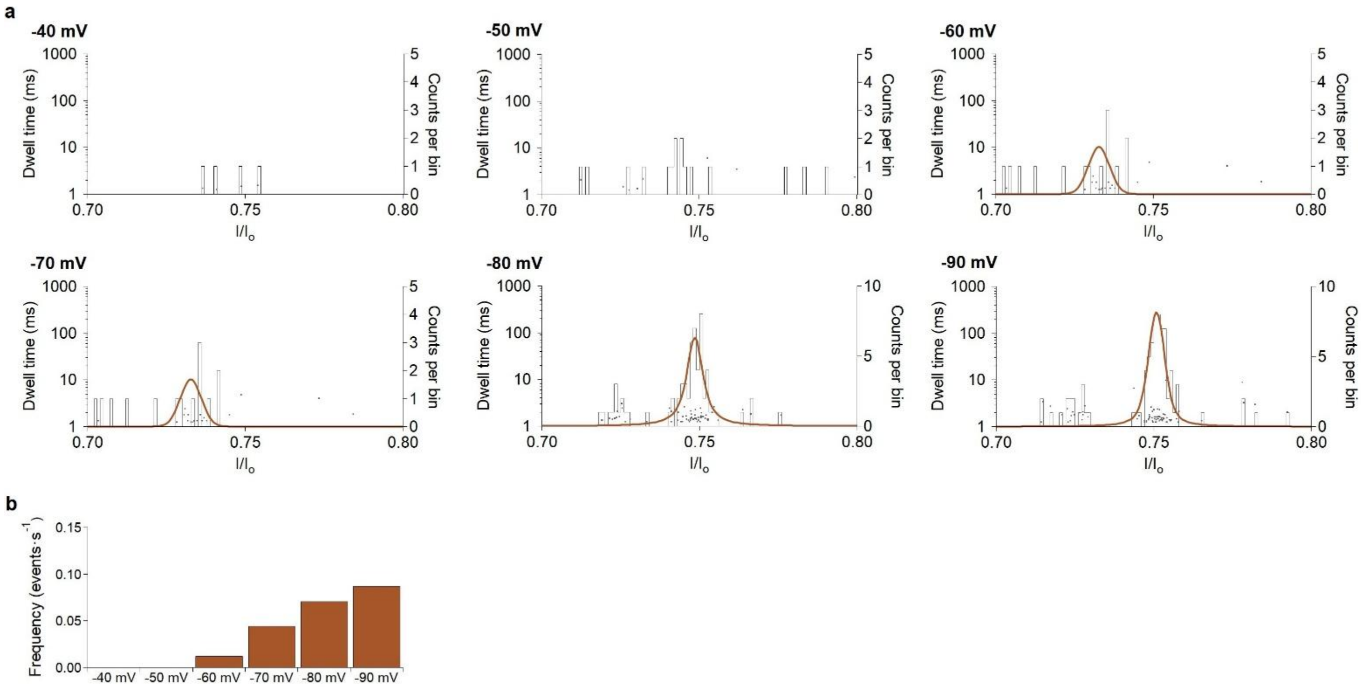


**SI Fig. 16: Nanopore sensing of synthetic MBP peptide [N91-R96].** Analysis of the peptide (final concentration: 5 µM) measured with AeL-R220S at applied potentials ranging from -40 mV to -90 mV. **a** Scatter plots (grey) of the dwell time versus the relative residual current (*I/I*$_o$). Voigt functions (colored) were fitted to histograms (black) of the corresponding *I/I*$_o$ distributions. No peak fitting was performed at -40 mV and -50 mV due to the low number of detected events. **b** Event frequencies of the blockade populations detected at different applied potentials.

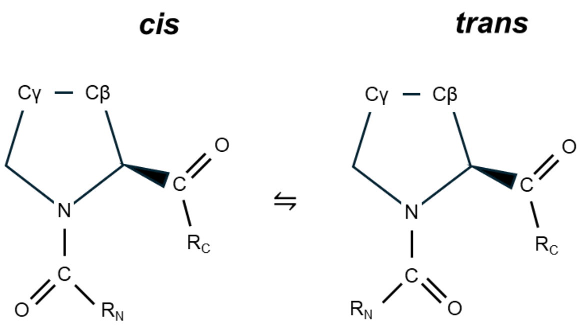


**SI Fig. 17: Schematic illustration of proline *cis* and *trans* conformation.**

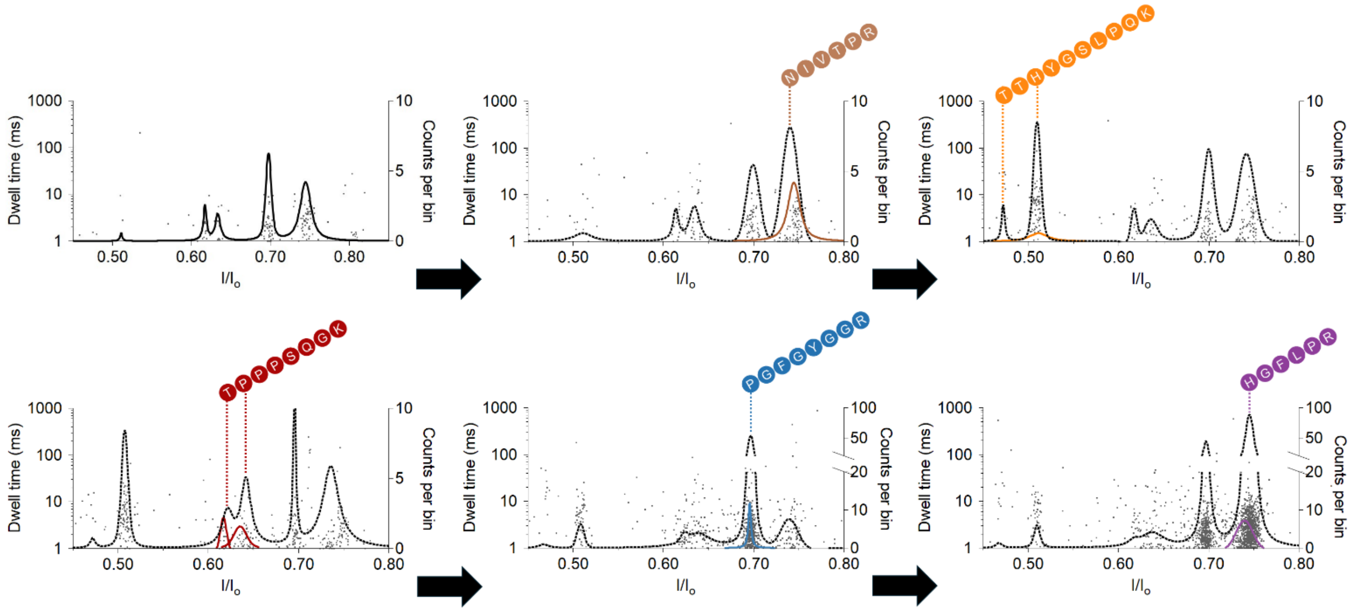


**SI Fig. 18: Nanopore sensing of an SP3-processed MBP digest with sequential spike-in of synthetic MBP peptides for peptide assignment.** Scatter plots (grey) of the dwell time versus the relative residual current ($I/I_o$) of the MBP digest (final concentration: 1 µM) and synthetic MBP peptides (final concentration: 5 µM) measured with AeL-R220S at -60 mV. Voigt functions (black / colored) were fitted to histograms of the corresponding $I/I_o$ distributions; the histograms are not shown for clarity. Synthetic MBP peptides were sequentially spiked into the MBP digest to validate the assignment of individual blockade populations. The solid lines indicate the corresponding peak populations in the unspiked MBP digest, whereas the dashed lines show the peak distributions in each measurement after addition of the corresponding synthetic peptide.

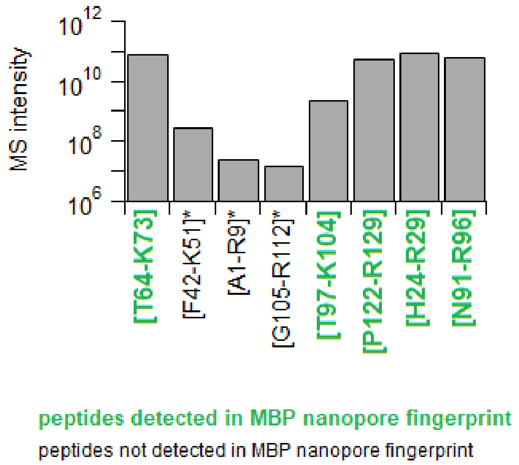


**SI Fig. 19: LC-MS/MS analysis of MBP-derived peptides detected in the digest and comparison with nanopore measurements.** Peptides highlighted in green were detected in the nanopore fingerprint, whereas the remaining peptides (*containing missed cleavages) were not detected in the nanopore measurements. These peptides generally exhibited lower LC-MS/MS signal intensities.
